# Lipid-anchored Proteasomes Control Membrane Protein Homeostasis

**DOI:** 10.1101/2023.05.12.540509

**Authors:** Ruizhu Zhang, Shuxian Pan, Suya Zheng, Qingqing Liao, Zhaodi Jiang, Dixian Wang, Xuemei Li, Ao Hu, Xinran Li, Yezhang Zhu, Xiaoqi Shen, Jing Lei, Siming Zhong, Xiaomei Zhang, Lingyun Huang, Xiaorong Wang, Lan Huang, Li Shen, Bao-Liang Song, Jingwei Zhao, Zhiping Wang, Bing Yang, Xing Guo

## Abstract

Protein degradation in eukaryotic cells is mainly carried out by the 26S proteasome, a macromolecular complex not only present in the cytosol and nucleus but also associated with various membranes. How proteasomes are anchored to the membrane and the biological meaning thereof have been largely unknown in higher organisms. Here we show that N-myristoylation of the Rpt2 subunit is a general mechanism for proteasome-membrane interaction. Loss of this modification in the Rpt2-G2A mutant cells leads to profound changes in the membrane-associated proteome, perturbs the endomembrane system and undermines critical cellular processes such as cell adhesion, endoplasmic reticulum-associated degradation (ERAD) and membrane protein trafficking. Rpt2^G2A/G2A^ homozygous mutation is embryonic lethal in mice and is sufficient to abolish tumor growth in a nude mice xenograft model. These findings have defined an evolutionarily conserved mechanism for maintaining membrane protein homeostasis and underscored the significance of compartmentalized protein degradation by <u>m</u>yristoyl-<u>a</u>nchored <u>p</u>roteasomes (MAPs) in health and disease.

## Introduction

Proteins in eukaryotic cells are highly compartmentalized, so are their synthesis and degradation (*1, 2*). Cells have evolved multiple strategies to dispatch proteins to different locations for degradation under normal or stressed conditions (*2–7*). This implies that the 26S proteasome, which digests the majority of eukaryotic proteins, must be available when substrates emerge or arrive. Consisting of a 20S core particle (CP) and one or two 19S regulatory particles (RP), the 26S proteasome is highly abundant but not evenly distributed in cells. How subcellular localization of these macromolecular complexes is determined is still not fully understood (*8–11*). In particular, a number of studies since early 1990’s have reported proteasomes that bound to various membrane structures of cells from different tissues and species (*11*). In some cases, this was attributed to proteasome binding to specific membrane proteins, whereas a general mechanism for proteasome-membrane association has been lacking, precluding further investigation of its biological meaning.

On the other hand, proteasomes are known to be regulated by a variety of chemical modifications including N-myristoylation (*12*), a co-translational lipid modification on proteins that start with methionine and glycine (Met_1_-Gly_2_). After removal of the initiator Met, the myristoyl group can be covalently conjugated to Gly2 by N-myristoyltransferase 1 and 2 (NMT1/2) (*13, 14*). In HeLa cells, scores of proteins were found to be modified this way, most of which are membrane-targeted (*15*). Secure membrane anchoring of myristoylated proteins requires additional modifications (e.g. palmitoylation) or electrostatic interaction between positively charged amino acids with the negatively charged head groups of phospholipids. In the latter case, membrane association can be antagonized by a phosphorylation event nearby (*16, 17*). Since no endogenous demyristoylating enzymes have been identified, N-myristoylation is currently considered irreversible. Nonetheless, the bacterial effector protein IpaJ from *Shigella* can specifically cleave after the myristoylated glycine residue and cause a proteolytic loss of protein myristoylation in the host cell (*18, 19*).

Rpt2/PSMC1, a component of the 19S RP, is the only proteasome subunit that begins with Met_1_-Gly_2_, thus the only subunit that can be N-myristoylated. Rpt2-Gly2 myristoylation has been detected in multiple mass spectrometry (MS) studies from yeast to plants to mammals (*15, 20–24*), and the N-terminal MG motif of Rpt2 is invariant through evolution (fig. S1A). In yeast, Rpt2 myristoylation has been suggested to target proteasomes to the nuclear envelope (*25*), and deletion or mutation of the myristoylation site (Rpt2-ΔG or G2A) led to impaired quality control of nuclear proteins (*26*). In human cells, Rpt2 appears to be one of the most myristoylated proteins, similar to the well-established membrane-localized kinases such as Src and PRKACA (*15*). However, the function of Rpt2 myristoylation in higher eukaryotes has not been investigated, except for our recent finding that it is required for Rpt2-Tyr439 phosphorylation by Src (*27*).

## Rpt2 myristoylation anchors proteasomes to the membrane in mammalian cells

We confirmed Rpt2 myristoylation in mammalian cells and its presence in the assembled proteasomes using two methods of detection. First, a clickable analog of myristic acid, Alk-Myr (*15*), was used to metabolically label 293T cells stably expressing TBHA-tagged α3, a 20S subunit (Fig. 1A). The TBHA tag contains a biotinylation sequence and a HA tag that facilitate the isolation and detection of proteasomes (*28*). Streptavidin-purified proteasome complexes from these cells were subjected to an *in vitro* click reaction, and myristoylated endogenous Rpt2 could be readily detected from the 26S proteasomes only in the presence of Alk-Myr (Fig. 1B and fig. S1B). Second, we developed a myristoylation-specific antibody against the N-terminus of Rpt2, which detected myristoylation of Rpt2 in mammalian cells but not unmodified Rpt2 recombinantly expressed from *E. coli* (fig. S1C). The immunoreactivity was completely eliminated by the G2A mutation (fig. S1D) or by overexpression of IpaJ (fig. S1E). Rpt2 myristoylation was also abrogated by IMP-1088, a specific inhibitor NMT1/2 (*29*) and greatly diminished by Tris (dibenzylideneacetone) dipalladium (Tris-DBA), a generic NMT1 inhibitor (*30*)(fig. S1F, G), further confirming the specificity of our detection methods. N-myristoylated Rpt2 was widely seen in total extracts from all cell/tissue types examined and on proteasome complexes affinity-purified from different organs of the Rpn11-TBHA knock-in mice (*31*) (Fig. 1C, D and fig. S1H, I). Sucrose gradient ultracentrifugation and native gel electrophoresis assays both showed that N-myristoylated Rpt2 was primarily present in fully assembled 26S/30S proteasomes (fig. S1J and fig. S3D), consistent with its co-precipitation with the 20S CP (Fig. 1B). Together, these characterizations demonstrate that Rpt2 myristoylation indeed occurs in mammalian cells and on the proteasome holoenzyme.

**Fig. 1.**
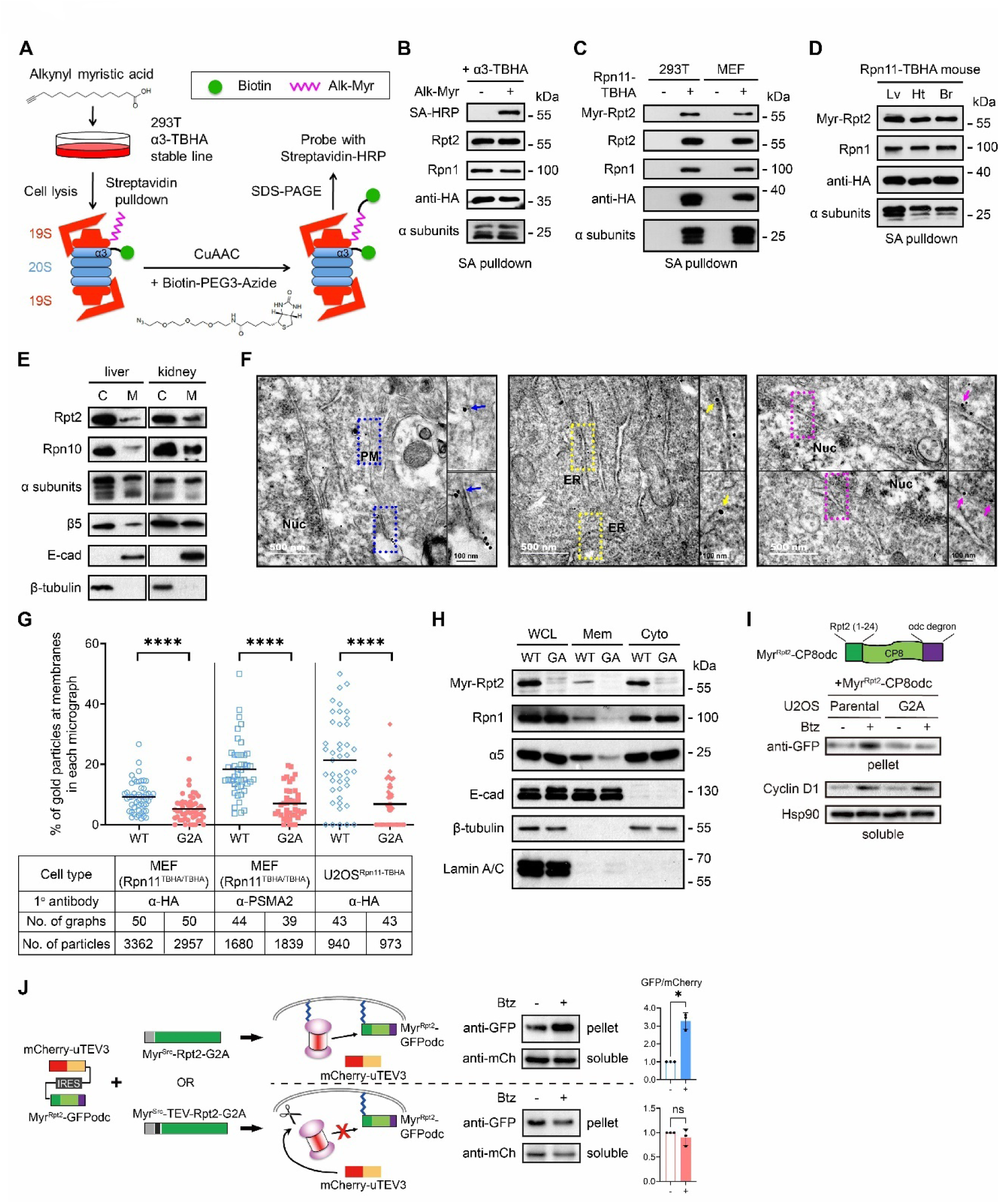
Rpt2 myristoylation mediates proteasome-membrane association. (**A**) Procedure of detecting endogenous Rpt2 myristoylation by metabolic labeling and click chemistry. CuAAC, copper(I)-catalyzed azide alkyne cycloaddition. (**B**) Proteasome samples purified by streptavidin (SA) pulldown were subjected to click chemistry reaction as in (A) and probed with SA-HRP and the indicated antibodies. (**C**) Western blot detection of Rpt2 myristoylation from proteasomes affinity-purified from 293T cells stably expressing Rpn11-TBHA or from Rpn11^TBHA/TBHA^ MEFs. 293T parental cells and WT MEFs were respectively used as control (“-”). (**D**) Immunoblot of myristoylated Rpt2 from proteasomes purified from the liver (Lv), heart (Ht) and brain (Br) tissues of Rpn11^TBHA^ mice. (**E**) Immunoblot of proteasome subunits in cytosolic (C) and membrane (M) fractions of mouse tissues. E-cadherin (E-cad) and β-tubulin mark the membrane and cytosolic fractions, respectively. (**F**) Representative IEM images of cerebral cortex slices from Rpn11^TBHA^ mice stained with an anti-HA antibody. Boxed areas are enlarged and shown on the right, with membrane-localized gold particles indicated by arrows of colors corresponding to different membranes. Nuc, nucleus; ER, endoplasmic reticulum; PM, plasma membrane. (**G**) Quantified results of IEM experiments performed with the indicated cell types and antibodies. Each data point represents an independent micrograph. Lines indicate the means of each group. ****, *P* < 0.0001 (Mann Whitney test, two-tailed). (**H**) Membrane (Mem) and cytosolic (Cyto) fractions as well as whole cell lysates (WCL) of WT and G2A (GA) MEFs were probed with the indicated antibodies. Lamin A/C was used as a nuclear protein marker. (**I**) Single clones of U2OS parental and G2A knock-in cells stably expressing the Myr^Rpt2^-CP8odc reporter (showon on the top) were treated with Bortezomib (Btz, 5 μM, 3 h). Membrane (pellet) and soluble fractions were separated and analyzed by immunoblotting. (**J**) (Left) A schematic representation of the cleavable Rpt2 system for assessing membrane-associated proteasome activity. After Btz treatment, the Myr^Rpt2^-GFPodc reporter was immunoblotted and quantified (right).

As we have seen before (*27*), N-myristoylation is essential for Rpt2 association with the membrane, since the Rpt2-G2A mutant was completely absent from membrane fractions of the cell (fig. S2A). Fusing the myristoylation sequence of Src (aa. 1-14) to the N-terminus of Rpt2-G2A (Myr^Src^-Rpt2-G2A) restored membrane association of the latter without affecting its assembly into the proteasome (fig. S2A, B), confirming the specific role of the myristoyl group in Rpt2 membrane anchoring. It should be noted, though, that like many myristoylated proteins with an N-terminal electrostatic switch, membrane-binding of WT Rpt2 is also dependent on a stretch of basic residues conserved through evolution (aa. 15-24, fig. S2A); and that this interaction may be disrupted by phosphorylation of Ser4 near the N-terminus (fig. S2A), which is one of the most frequently detected proteasome phosphosites conserved in vertebrates (*31*).

To determine whether Rpt2 myristoylation can mediate membrane binding of the proteasome complex, we first confirmed by mouse tissue fractionation that, in addition to Rpt2, other RP and CP subunits of the proteasome were also present in the membrane fractions (Fig. 1E). We then took advantage of our Rpn11-TBHA knock-in mice and optimized an immunogold electron microscopy (IEM) protocol to detect endogenous 26S proteasomes with anti-HA antibodies in cerebral cortex sections. The results further demonstrated the presence of proteasomes at not only the plasma membrane but also the nuclear envelope, endoplasmic reticulum (ER) and mitochondria of neurons (Fig. 1F and fig. S2C).

Next, we obtained Rpt2^+/+^;Rpn11^TBHA/TBHA^ and Rpt2^G2A/G2A^;Rpn11^TBHA/TBHA^ mouse embryonic fibroblasts (MEFs) by crossing Rpn11^TBHA/+^ mice with Rpt2^G2A/+^ mice. IEM with anti-HA and anti-PSMA2 (a 20S subunit) antibodies on WT MEFs (Rpt2^+/+^;Rpn11^TBHA/TBHA^) showed proteasome localization at additional membranes including the endosome, lysosome and the Golgi apparatus (fig. S2C), consistent with several earlier reports (*32–34*). Overall, approximately 5-10% of gold particles were found to be membrane-localized from all the micrographs that contained evident membranous structures. Importantly, membrane association of the proteasome was significantly reduced in the Rpt2^G2A/G2A^ MEFs, and a similar trend was observed in U2OS^G2A/G2A^ cells generated by CRISPR/Cas9 (Fig. 1G). Further analysis of the IEM results revealed that loss of Rpt2 myristoylation mainly interfered with proteasome localization at the plasma membrane, ER, Golgi apparatus and endosomes/lysosomes (“PERGEL”), whereas mitochondria- and nuclear envelope-associated proteasomes (“MN”) were less affected (fig. S2D). This pattern agrees well with the global distribution of myristoylated proteins identified by proteomics (*15*) and suggests that Rpt2 myristoylation can direct the proteasome to various subcellular locations. Moreover, cell fractionation assays also demonstrated a clear decrease of 19S and 20S components in the membrane fraction from Rpt2-G2A cells compared to control (Fig. 1H). Finally, using total internal reflection fluorescence imaging with structured illumination microscopy (TIRF-SIM), we observed distinct signals of GFP-tagged Rpn10 (another 19S subunit) in the close vicinity of ventral plasma membrane in WT cells, but much less so in G2A knock-in cells (fig. S2E). Altogether, these results indicate that Rpt2 N-myristoylation is a major and prevalent mechanism mediating proteasome-membrane association. We hereby refer to these <u>m</u>yristoyl-<u>a</u>nchored <u>p</u>roteasomes as MAPs.

To specifically detect the activity of MAPs toward membrane-bound protein substrates, we fused the Rpt2 N-terminal sequence (aa. 1-24 including the myristoylation site and the polybasic region)(*27*) to a commonly used model substrate of the proteasome, GFPodc (*35*) or its loosely folded variant CP8odc (*36*)(Fig. 1I and fig. S2F). The resulting membrane-targeted Myr^Rpt2^-CP8odc reporter was stabilized by the proteasome inhibitor Bortezomib (Btz) in the membrane fraction of WT cells, suggesting that it underwent constant local proteasomal degradation (Fig. 1I). However, membrane-bound Myr^Rpt2^-CP8odc showed no response to Btz in the G2A cells (Fig. 1I), consistent with the lack of proteasome activity at the membrane and also implying that cytosolic proteasomes in G2A cells cannot efficiently degrade this membrane-localized substrate. Btz-induced accumulation of non-membrane substrates (e.g., Cyclin D1) was not affected by the G2A mutation (Fig. 1I). As these results were obtained from single clones of cells stably expressing the reporter protein, we wanted further confirmation by acutely disrupting proteasome-membrane interaction in a cell population. To this end, a cleavable version of Myr^Src^-Rpt2-G2A was engineered by inserting the tobacco etch virus (TEV) protease recognition motif (ENLYFQ|S) behind the Myr^Src^ sequence. The resulting Myr^Src^-TEV-Rpt2-G2A could associate with the membrane just like Myr^Src^-Rpt2-G2A but could be cut and released from the membrane upon TEV protease expression (fig. S2A, B and G). We knocked down endogenous Rpt2 in WT cells, simultaneously expressed either of these two forms of Rpt2, then co-introduced Myr^Rpt2^-GFPodc together with uTEV3, an optimized version of the TEV protease (*37*)(Fig. 1J). Bortezomib treatment of cells expressing Myr^Src^-Rpt2-G2A (non-cleavable) led to accumulation of Myr^Rpt2^-GFPodc as expected. In contrast, this reporter was insensitive to Btz treatment in cells expressing Myr^Src^-TEV-Rpt2-G2A, in which the Rpt2 protein had been dissociated from the membrane by uTEV3 (Fig. 1J). This latter result is very similar to that in G2A knock-in cells (Fig. 1I). Such acute perturbation of membrane binding further demonstrates the direct contribution of MAPs to compartmentalized protein degradation at the membrane.

Despite the local effect, loss of Rpt2 myristoylation did not affect overall proteasomal abundance, activity or cell viability (fig. S3A-C). Proteasome holoenzyme assembly remained intact in Rpt2-G2A cells as judged by native gel electrophoresis (fig. S3D) and pulldown-MS experiments (fig. S3E). However, dislodgement of the G2A proteasome from the membrane considerably altered its interactome. Nearly 500 proteasome interacting proteins (PIPs)(*23*) were identified in both WT and G2A cells from two independent MS experiments, over a third of which showed enhanced association with the G2A proteasome (fig. S3F). These proteins are located throughout the cell, mostly in the cytoplasm and nucleus (fig. S3F). On the other hand, a few PIPs exhibited reduced proteasome binding in the G2A cells. Interestingly, these included the membrane-localized tyrosine phosphatase PTPN2 and tyrosine kinase Src, which together regulate Rpt2-Y439 phosphorylation and membrane proteasome function as we recently reported (*27*)(fig. S3E, F). Also reduced/lost in the G2A proteasome interactome was the non-essential proteasome chaperone, Ecm29 (fig. S3F), which is known to locate at membrane-bound organelles such as endosomes and the ER (*32, 38*). Therefore, blocking Rpt2 myristoylation may re-partition proteasomes between the membrane and soluble compartments, potentially leading to proteomic and functional alterations of the cell.

## MAPs are required for proper turnover and processing of membrane-related proteins

To gain insights into the proteins selectively controlled by MAPs, we performed quantitative mass spectrometry on total cell extracts of WT and G2A MEFs with reciprocal SILAC labeling (stable isotope labeling using amino acids in cell culture). A total of 6774 proteins were detected from three biological repeats, with 4470 proteins identified in all 3 runs (Fig. 2A, fig. S4A). Nearly one tenth of these proteins (436) were differentially regulated in G2A cells (|Log_2_FoldChange(G2A/WT)| > 0.585), with 66.6% of them being membrane-/organelle-associated based on annotations from UniProt and the Human Protein Atlas (Fig. 2B). These include peripheral membrane proteins, transmembrane proteins and luminal/secreted proteins, which are widely localized at various subcellular sites and involved in a variety of biological functions (Fig. 2B, C, fig. S4B, C). Overall, there was minimal correlation between their protein and transcript levels (*r*^2^ = 0.099), agreeing with a post-translational mechanism underlying their differential expression (fig. S4D). Of note, the majority of these proteins showed increased abundance within G2A cells (Fig. 2B, C and fig. S4B), and the upregulation of multiple transmembrane and peripheral proteins was further confirmed by immunoblotting (Fig. 2D and fig. S4E). In addition to membrane localization, these upregulated proteins also shared the following features: 1) they became accumulated in WT cells following Btz treatment; 2) in G2A cells, however, they remained high and exhibited no or only minor responses to Btz (Fig. 2D); 3) as noted above, the mRNA expression of most of these proteins was comparable between WT and G2A cells (fig. S4E); 4) their elevated protein levels could be reversed by expressing the membrane-targeted Myr^Src^-Rpt2-G2A in the G2A cells (fig. S4F). These data indicate that MAPs are uniquely responsible for the turnover of a subset of proteins with diverse subcellular localizations and functions, and such turnover is largely blocked in cells lacking Rpt2 myristoylation.

**Fig. 2.**
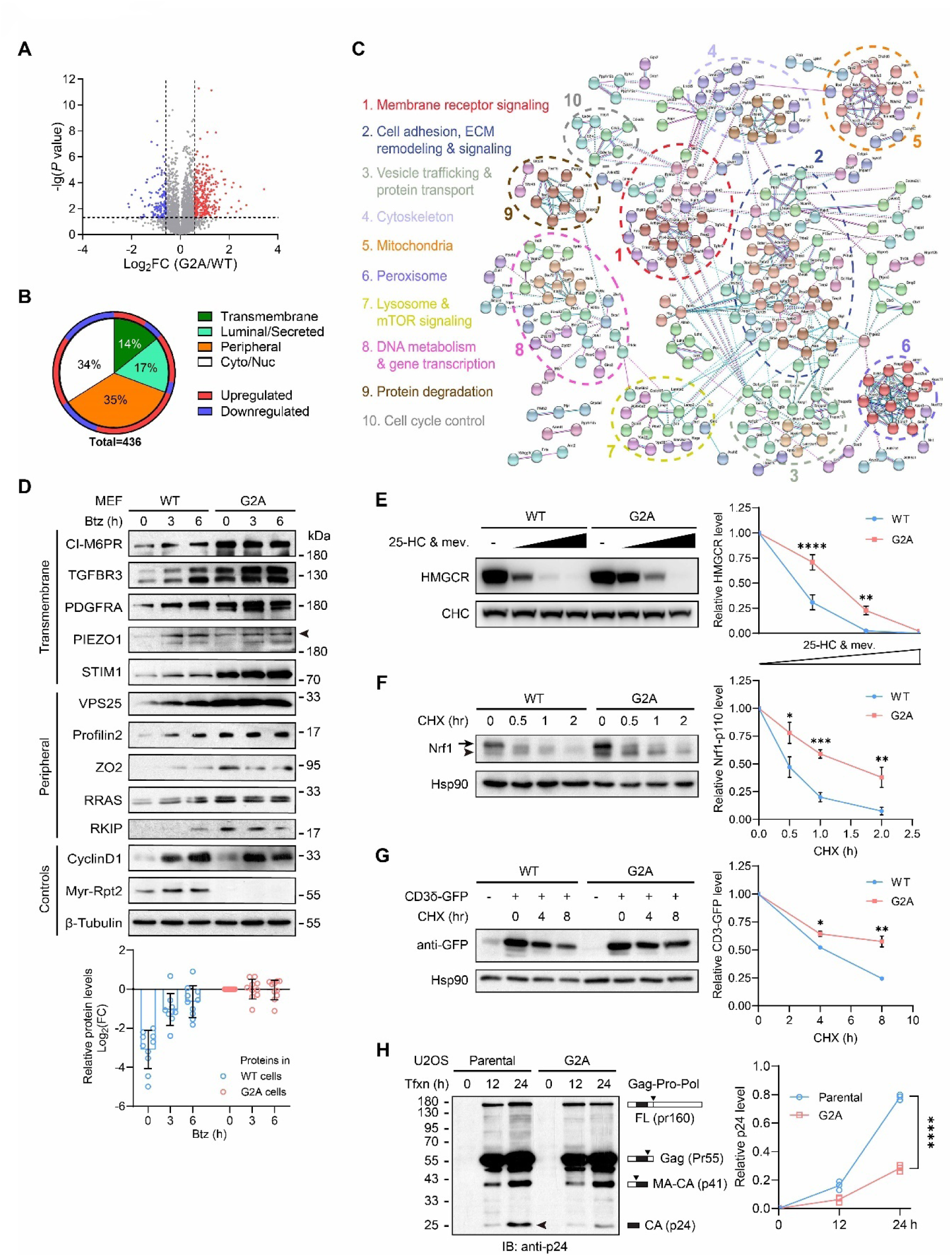
Loss of myristoylation interferes with membrane protein turnover and processing. (**A**) Volcano plot of SILAC-MS studies with WT and G2A MEFs. (**B**) Summary of differentially regulated proteins in G2A cells and their subcellular locations. (**C**) Protein-protein interaction network of upregulated proteins in G2A MEFs showing their subcellular distribution and biological functions. (**D**) WT and G2A MEFs were treated with Btz (1 μM) for the indicated time and probed for representative transmembrane and peripheral proteins. Arrowhead points to the PIEZO1 band (292 kDa). (Bottom) The relative abundance of each protein from these blots (represented by individual circles) was quantified after normalization to their basal levels in G2A cells. (**E**) WT or G2A MEFs were treated with increasing concentrations of 25-hydroxycholesterol (25-HC) and mevalonate (mev) for 4 hours. Denatured lysates were probed with the indicated antibodies. CHC (clathrin heavy chain) was shown as loading control. (**F**) Cells were treated with cycloheximide (CHX, 50 μg/ml) for the indicated time and turnover of endogenous Nrf1 was determined. Arrow, full-length Nrf1. Arrowhead, the p110 fragment. (**G**) Immunoblot of CD3δ-GFP after treatment with CHX. (**H**) Cells were transfected with the psPAX2 plasmid encoding the full-length (FL) Gag-Pro-Pol precursor protein. At the indicated time after transfection (tfxn), cells were harvested and probed with an anti-p24 antibody that recognizes the different protein fragments depicted. Downward triangles mark the major cleavage sites. MA, matrix protein. CA, capsid protein. The final product (CA/p24) is indicated with an arrowhead, and its relative level was normalized to the amount of Pr55 at each time point. All immunoblots above were repeated at least 3 times. *, *P* < 0.05; **, *P* < 0.01; ***, *P* < 0.001; ****, *P* < 0.0001 (Student’s *t*-test, n = 3).

An important mechanism for membrane protein regulation and quality control is through ER-associated degradation (ERAD), the end point of which is believed to be proteasomes in the cytosol (*39*). We assessed the turnover of three well-recognized ERAD substrates in WT and G2A cells, all of which are transmembrane proteins. First, HMGCR (3-hydroxy-3-methylglutaryl-coenzyme A reductase), the rate-limiting enzyme in cholesterol synthesis, can be rapidly degraded via ERAD (*40*). Sterol stimulation triggered HMGCR degradation in a dose-dependent manner in WT MEFs (Fig. 2E). In G2A MEFs, however, this effect was clearly weakened (Fig. 2E). Similarly, degradation of the ER-resident transcription factor Nrf1/NFE2L1 was also less efficient in G2A cells, resulting in a prolonged presence of the truncated p110 fragment of Nrf1/NFE2L1 (*41, 42*) (Fig. 2F). The third example was CD3δ-GFP. When not assembled into the T cell receptor complex, ectopically expressed CD3δ is known to be cleared by ERAD (*43*), which again occurred more slowly in G2A cells (Fig. 2G). All these findings highlight previously unknown contributions of the MAPs to efficient ERAD. Consistent with this notion, a recent cryo-electron tomography study visualized 26S proteasome clusters on the ER membrane engaging in ERAD (*44*). These proteasomes could well be anchored via Rpt2 myristoylation, which allows for the proteasome to meet the substrates as they exit ER and facilitates rapid degradation on the spot.

We also examined a pathologically relevant protein lipid-anchored to the plasma membrane, the Gag protein of human immunodeficiency virus (HIV). It is well established that N-myristoylation and plasma membrane association of Gag is critical for its processing, viral particle assembly and virus budding (*45, 46*). Proteolysis and maturation of Gag from HIV and multiple other retroviruses also depend on proteasome activity (*47, 48*). We expressed the full-length Gag-Pro-Pol protein in U2OS cells and monitored the processing of Gag with time. The 55 kDa Gag protein emerged similarly in control and G2A cells, but subsequent production of the p24 fragment (capsid protein, CA), which depends on membrane binding, was much delayed in U2OS-G2A cells (Fig. 2H). This effect echoes previous results obtained with proteasome inhibitors (*47*) and suggests a specific requirement for MAPs in viral protein processing.

## Dysregulation of endomembrane homeostasis and protein trafficking in cells lacking MAPs

Also differentially expressed in G2A cells were numerous proteins implicated in organellar function and intracellular trafficking (Fig. 2C). For instance, cation-independent mannose-6-phosphate receptor (CI-M6PR, also known as Igf2r), a key mediator of protein sorting between Golgi and lysosomes/late endosomes, and its Golgi-resident receptor Golga4/Golgin-245 (*49, 50*) were both upregulated in G2A MEFs and could be returned to normal levels upon Myr^Src^-Rpt2-G2A expression (Figs. 2C, 2D, 3A, fig. S4F). In addition, vacuolar protein-sorting-associated protein 25 (VPS25) and its binding partner VPS36 were both elevated in G2A cells (Fig. 2C, D). These proteins are central components of the ESCRT-II complex (endosomal sorting complex required for transport II) required for sorting of endosomal cargo proteins into multivesicular bodies (MVBs) then lysosomes (*51*). These findings prompted us to examine the relevant compartments. Compared to WT control, G2A cells exhibited significantly stronger LysoTracker staining with concurrent increase of several lysosomal proteins (Fig. 3B, C). Transmission electron microscopy (TEM) also showed an increase in the number of lysosomes/MVBs in G2A cells, whereas their sizes were comparable to those in WT cells (fig. S5A). Nonetheless, we occasionally observed extraordinarily large, membrane-bound vacuoles/inclusions in G2A cells that were not seen in WT cells (fig. S5A). On the other hand, Golgi cisternae appeared swollen in G2A cells as revealed by TEM, with a significant increase in the mean luminal width of individual cisterna as compared to control (Fig. 3D). The G2A MEFs were also more sensitive to the Golgi stress inducer, monensin (*52*)(Fig. 3E), indicating impairment of Golgi function.

**Fig. 3.**
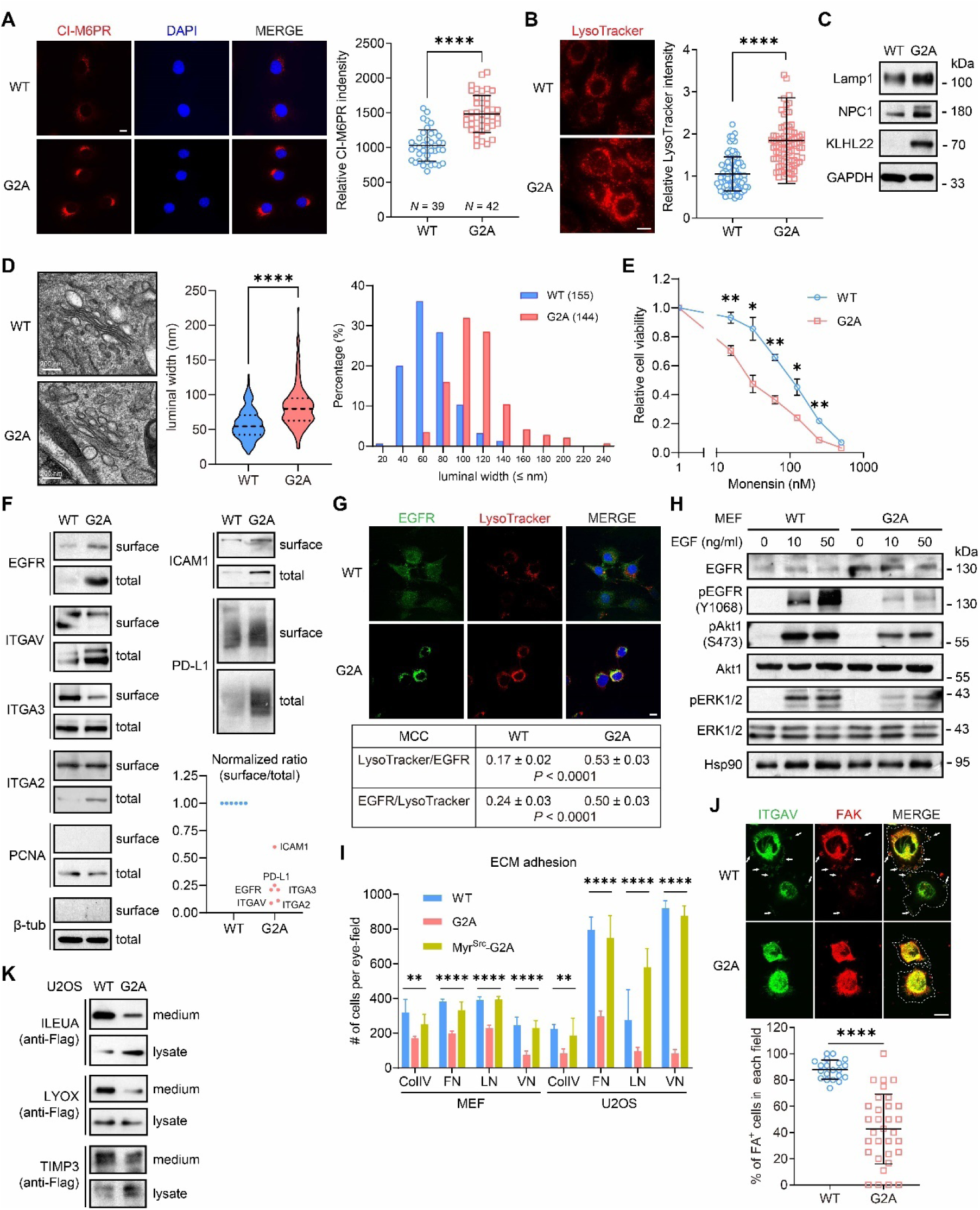
Perturbation of the endomembrane system and protein transport in Rpt2-G2A cells. (**A**) Immunofluorescence staining of CI-M6PR in G2A MEFs. Scale bar = 10 μm. ****, *P* < 0.0001 (two-tailed *t*-test, unpaired). (**B**) LysoTracker staining of MEFs. Scale bar = 10 μm. ****, *P* < 0.0001 (two-tailed *t*-test, unpaired. *N* = 88 each). (**C**) Western blot analysis of lysosome-related proteins in MEFs. (**D**) Representative TEM images of the Golgi apparatus in WT and G2A MEFs. (E) Luminal width of individual Golgi cisterna was measured (*60*) and quantified on the right. ****, *P* < 0.0001 (Welch’s *t*-test). (**E**) WT and G2A MEFs were treated with increasing concentrations of monensin for 48 h and released for another 24 h. Cell viability was measured. *, *P* < 0.05; **, *P* < 0.01 (two-tailed *t*-test, paired, *N* = 3). (**F**) Cell surface biotinylation assay of the indicated proteins. Each blot is representative of at least 3 independent experiments. The surface/total ratio of each protein was normalized to that in WT cells. EGFR, integrins and PCNA were from MEFs. ICAM1, PD-L1 and β-tubulin were from U2OS cells. PCNA and β-tubulin were shown as controls for the specificity of surface labeling. (**G**) Co-staining of MEFs with anti-EGFR antibody and LysoTracker. Scale bar = 10 μm. Mander’s colocalization coefficients (MCC) between the two channels were calculated for both cell types and presented as mean ± SEM. *P* values were determined by two-tailed *t*-test (unpaired). WT, *N* = 72; G2A, *N* = 76. (**H**) MEFs were serum-starved for 24 hours and treated with different concentrations of EGF for 5 min. Total cell lysates were blotted. (**I**) Quantified results of cell adhesion assays with WT and G2A MEFs on surfaces coated with different ECM proteins. ColIV, collagen IV; FN, fibronectin; LN, laminin; VN, vitronectin. **, *P* < 0.01; ****, *P* < 0.0001 (*N* = 18 eye-fields from 3 independent experiments, One-way ANOVA). (**J**) WT and G2A MEFs were seeded on fibronectin-coated coverslips for 30 min and immnostained with the indicated antibodies. Dotted lines mark cell contours. Arrows indicate peripheral focal adhesion (FA)-like structures (double-positive for ITGAV and FAK). Scale bar = 10 μm. ****, *P* < 0.0001 (two-tailed *t*-test, unpaired). WT, *N* = 20 eye-fields (318 cells); G2A, *N* = 32 eye-fields (179 cells). (**K**) C-terminally Flag-tagged expression constructs of the indicated secreted proteins were transfected into U2OS cells. Cell media were immunoprecipitated with anti-Flag antibody and probed together with each corresponding whole cell lysate.

To understand if such perturbation of the endomembrane system could lead to defects in protein trafficking, we examined the cell surface expression of several classic transmembrane proteins including EGFR, ICAM-1, PD-L1 and integrins (ITGA2/3/V) by surface biotinylation assays. Most of these proteins showed a higher total level in G2A cells. In contrast, all of them had a much lower surface/total ratio, meaning that a larger proportion of these proteins failed to reach or maintain at the plasma membrane in the absence of MAPs (Fig. 3F). Indeed, immunofluorescence staining demonstrated that the majority of EGFR in G2A MEFs was inside the cells, with a significantly higher fraction co-localizing with lysosomes than observed in WT cells (Fig. 3G). Functionally, the G2A cells responded poorly to EGF despite the higher total level of EGFR, and downstream activation of Akt and Erk1/2 were markedly undermined (Fig. 3H).

Reduced surface presentation of integrin molecules predicts weakened cell adhesion, which is consistent with our proteomic and transcriptomic analyses (Fig. 2C and figs. S4C, G) and our routine observation that G2A knock-in cells were easier to dissociate from the vessels by trypsinization than their WT counterparts (fig. S5B, C). Using defined extracellular matrix (ECM) components, we found that attachment of G2A cells to collagen IV-, fibronectin-, laminin- or vitronectin-coated surfaces was all significantly weaker than control, in agreement with attenuation of integrin function (Fig. 3I and fig. S5D). Upon seeding onto ECM-coated surface, WT cells readily spread out and formed focal adhesion-like structures positively stained with FAK and ITGAV at cell periphery, while this occurred much less efficiently in G2A cells (Fig. 3J). Importantly, the weakened adhesion of G2A cells could be rescued by expression of Myr^Src^-Rpt2-G2A (Fig. 3I and fig. S5D), strongly suggesting that the adhesion defect of G2A cells indeed resulted from the lack of MAPs. As expected, cell migration was also severely undermined by the Rpt2-G2A mutation in both wound-healing/scratch assays and transwell migration assays (fig. S5E, F).

Furthermore, secretion of certain extracellular proteins was also impeded by the G2A mutation (Fig. 3K). Such intracellular entrapment may explain their apparent “upregulation” in G2A MEFs as identified by mass spectrometry (Fig. 2C). Together, these data demonstrate that MAPs play a fundamental role in maintaining the homeostasis of the endomembrane system, thereby having a broad impact on protein trafficking along the secretory pathway.

## MAPs are essential for tumorigenesis and embryonic development

Membrane receptor signaling, cell adhesion, protein secretion and vesicular trafficking are all instrumental for oncogenesis. The ESCRT-II complex and CI-M6PR have also been suggested to be tumor suppressors (*49, 53*). To demonstrate the consequences of the above cellular changes *in vivo*, we adopted a xenograft tumor model. The immortalized MEFs (WT and G2A) were further transformed with hTERT/N-Ras^G12V^ (*54*) and implanted subcutaneously into nude mice (fig. S6A). Oncogene-transformed WT cells readily formed tumors as expected. In sharp contrast, the MEFs of G2A origin completely failed in tumorigenic growth (Fig. 4A, B). In tissue culture, however, the transformed G2A MEFs showed no growth defects (in fact they grew slightly better than WT MEFs, fig. S6B). This indicates that the inability of the G2A cells to grow as xenograft was probably due to anomalous interactions with their microenvironment *in vivo*, as can be deduced from the *in vitro* studies above. Again, replacement of endogenous Rpt2-G2A with Myr^Src^-Rpt2-G2A in transformed G2A MEFs largely restored tumorigenicity of the cells (Fig. 4A, B), highlighting the critical role of MAPs in tumorigenesis.

**Fig. 4.**
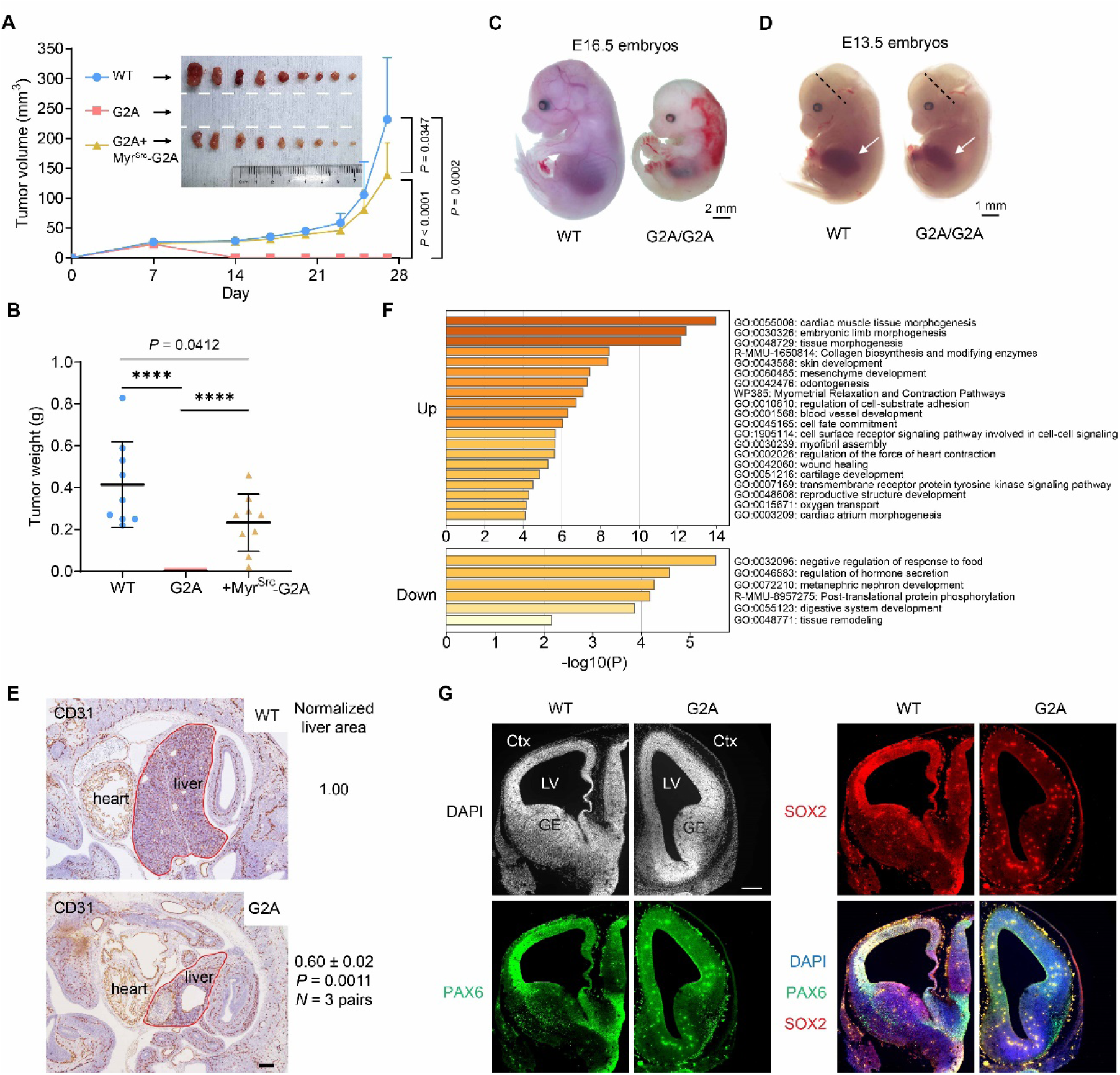
Proteasome myristoylation is required for tumorigenesis and embryonic development. (**A**) and (**B**) The indicated transformed MEFs were subcutaneously grafted into nude mice, and the sizes of xenograft tumors were monitored (A). Dissected tumors were photographed (inset of A) and weighed (B). ****, *P* < 0.0001, Welch’s *t*-test, *N* = 9. (**C**) and (**D**) Representative images of WT and G2A/G2A mouse embryos (littermates) at E16.5 (C) and E13.5 (D). Arrows point to the fetal liver. Dotted lines indicate positions of the coronal sections shown in (G). (**E**) Immunohistochemistry staining of E13.5 embryos. Scale bar = 200 μm. Outlines of the the liver area are marked. Normalized liver sizes are shown on the right (twp-tail paired *t*-test). Anti-CD31 stains blood vessels. (**F**) Gene Ontology (GO) term analysis of differentially expressed mRNAs in G2A fetal livers as determined by RNA-Seq. (**G**) Immunohistochemistry staining of E13.5 mouse brain. Corresponding coronal sections of WT and G2A embryos (as shown in D) are juxtaposed for direct comparison. LV, lateral ventricle; GE, ganglionic eminence; Ctx, cortex. Scale bar = 200 μm.

Moreover, homozygous Rpt2-G2A mutation in mice caused embryonic lethality with 100% penetrance (Fig. 4C, D). The G2A/G2A embryos died before E16.5, showing hemorrhage at the dorsal and interdigital regions with other parts of the body being anemic (Fig. 4C). In mutant embryos at earlier stages (e.g. E13.5), severe developmental defects of vital organs such as liver and brain could be detected by histology (Fig. 4D-G). Not only the fetal liver was much smaller in the G2A embryos, but the lobule-like organization of liver cells was also largely absent (Fig. 4E). Transcriptomic analysis of mutant fetal livers identified a small fraction of genes with aberrant expression, which are enriched in essential development-related activities including cell adhesion, tissue morphogenesis and cell fate determination (Fig. 4F). The mutant brain also exhibited abnormal morphology. PAX6^+^ and SOX2^+^ neural progenitor/stem cells were greatly reduced in number and showed distinct distributions from those in WT embryos, indicative of delayed neurogenesis and impaired cell migration (Fig. 4G). In fact, the highest level of Rpt2 myristoylation in the brain is seen at the pre- and peri-natal stages, mirroring its indispensable role during brain development (fig. S6C). Together, these phenotypes provide strong *in vivo* evidence for the physiological significance of MAPs in mammals, which is well in line with the strict conservation of Rpt2 myristoylation from yeast to human.

## Discussion

Among several mechanisms through which the proteasome can be targeted to the membrane (*11*), our results favor Rpt2 myristoylation as a major determinant of proteasome-membrane interaction. This has allowed us to interrogate the biological meaning of membrane-localized proteasomes by studying the Rpt2-G2A mutant. Albeit a minority of the entire proteasome pool in cells, MAPs apparently play irreplaceable roles at both cellular and organismal levels, consistent with their wide presence at various subcellular locales, in many different cell types and at different developmental stages. This may be attributed to the co-translational and irreversible nature of Rpt2 N-myristoylation, although its level does change in the developing brain (fig. S6C) for reasons unknown. One would therefore expect that the amount of membrane-associated proteasomes, which may also be regulated by Rpt2-Ser4 phosphorylation and PIPs, can vary among cell types and in response to pathophysiological signals. Further studies on the dynamic regulation of proteasome-membrane interaction would be important for better understanding of compartmentalized degradation of membrane-related proteins.

Membrane anchoring of the proteasome is expected to increase its local concentration and/or facilitate substrate capture, as has been demonstrated for receptor tyrosine kinases (*55*). On one hand, the local presence of proteasomes can be particularly convenient for membrane-related events such as ERAD and Gag cleavage, even though these processes have been thought to be controlled solely by cytosolic proteasomes. On the other hand, in light of the exceedingly crowded cellular environment, membranes may serve as a foothold for myristoylated proteasomes and a platform for degradation of non-membrane-associated proteins, as we also noticed from the proteomic results (Fig. 2B). *In vivo*, e.g., during neurodevelopment, the membranous compartments of neuronal cells undergo drastic remodeling, and local protein synthesis and degradation are tightly coupled in membrane-enclosed structures such as spines and synapses (*1, 56–59*). This imposes heavy demands on the proteasome system including MAPs, the lack of which can lead to the brain development defects we observed in the G2A/G2A mouse embryos.

Besides mis-accumulation of substrates, elimination of MAPs also indirectly influences protein trafficking and secretion, which could be both cause and consequence of dysregulation of the endomembrane system of the cell. Although the aberrant upregulation of sorting proteins such as VPS25 and CI-M6PR in G2A MEFs are likely involved, more detailed mechanisms would require further investigation and the exact mediators are probably cell type- and organelle-specific. Some of these changes in G2A cells could in turn cause alterations of signal transduction and gene expression as revealed by our transcriptomic analysis. In this sense, compartmentalized protein regulation by MAPs has far-reaching biological significance, as supported by the embryonic lethality and potent tumor-suppressive effect of the Rpt2^G2A/G2A^ mutation. From a therapeutic viewpoint, selective targeting of MAPs while leaving the bulk of cellular proteasomes untouched may be effective in suppressing tumor growth with much reduced side effects associated with common proteasome inhibitors.

## Acknowledgement

We appreciate valuable inputs from Shixian Lin, Liangyi Chen, Hai Song, Hengyu Fan, Luyang Yu, James Zhijian Chen, Junfang Ji and past members of the Guo lab. We thank Yanfen Liu, Liang Shi, Shengda Lin, Bin Zhao and HuaBio for kindly providing crucial DNA plasmids, antibodies and reagents. We thank Guangzhou Computational Super-resolution Biotech Co., Ltd for their commercial super-resolution microscope (HIS-SIM) and assistance on data acquisition, image reconstruction and analysis. We are grateful for Qirou Wu, Qingzhe Wu, the Core Facility of Life Sciences Institute (Zhejiang University) and Hexia Luo (National Institute of Biological Sciences) for excellent technical assistance. This work was supported by grants from National Natural Science Foundation of China 31870762, 32071257, 32271303 (X.G.), 22074132 (B.Y.) 32022023 (L.S.) and from Zhejiang Natural Science Foundation LR18C050001, LZ23C050003 (X.G.) and LR20B050001 (B.Y.); National Institutes of Health grants R35GM145249 and R01GM074830 to L.H..

