## Supplemental Information for "Lipid-anchored Proteasomes Control Membrane Protein Homeostasis"

### Supplementary Materials

### Materials and Methods

#### Antibodies and reagents

The rabbit anti-myr-Rpt2 polyclonal antibody was generated with the peptide antigen (Myr)GQSQSGGHGPGGGKKD-Cys, followed by negative absorption and Protein A affinity purification. Informatin of all the other commercial antibodies and reagents is listed below.

| Antibodies | Source | Catalog No. and RRID | Application |
|---|---|---|---|
| 20S α123567 | Enzo Life Sciences | Cat# BML-PW8195; RRID: AB_11177877 | 1:5,000 (WB) |
| AKT1 | HUABIO | Cat# EM40507 | 1:1,000 (WB) |
| AKT1 (pS473) | HUABIO | Cat# ET1607-73 | 1:1,000 (WB) |
| CD31 | Cell Signaling | Cat# 77699S; RRID: AB_2722705 | 1:200 (IHC) |
| CI-M6PR | HUABIO | Cat# ET1602-5 | 1:2,000 (WB)<br>1:300 (ICC) |
| Clathrin Heavy Chain (CHC) | BD Biosciences | Cat# 610500; RRID: AB_397866 | 1:1,000 (WB) |
| Cyclin D1 | HUABIO | Cat# ET1601-31 | 1:1,000 (WB) |
| E-cadherin | Affinity Biosciences | Cat# BF0219; RRID: AB_2833860 | 1:1,000 (WB) |
| EGFR | Cell Signaling | Cat# 4267T; RRID: AB_2799342 | 1:1,000 (WB)<br>1:250 (ICC) |
| EGFR (Tyr1068) | Cell Signaling | Cat# 3777T; RRID: AB_1903957 | 1:1,000 (WB) |
| Erk (pY204) | Santa Cruz Biotechnology | Cat# sc-7383; RRID: AB_627545 | 1:500 (WB) |
| Erk1/2 | HUABIO | Cat# ET1601-29 | 1:1,000 (WB) |
| Flag-HRP | Shanghai Genomics Technology | Cat# GNI4310-FG; RRID: AB_2885081 | 1:5,000 (WB) |
| Flag-Tag | AbMart | Cat# M20008S; RRID: AB_2713960 | 1:5,000 (WB) |
| GAG | HUABIO | Cat# ER50102 | 1:1,000 (WB) |
| GAPDH | Millipore | Cat# CB1001; RRID: AB_2107426 | 1:1,000 (WB) |
| GFP | AbMart | Cat# M20004L; RRID: AB_2619674 | 1:1,000 (WB) |
| GOLGA4 | ABclonal | Cat# A10216; RRID: | 1:1,000 (WB) |

| | | | |
|---|---|---|---|
| | | AB_2757739 | |
| HA-Tag | Cell Signaling | Cat# 3724S; RRID: AB_1549585 | 1:5,000 (WB) |
| HMGCR | Produced from hybridoma (Baoliang Song) | IgG-A9 | 1:1,000 (WB) |
| Hsp90α | HUABIO | Cat# M1603-3 | 1:5,000 (WB) |
| ICAM1 | HUABIO | Cat# ET1609-46 | 1:1,000 (WB) |
| ITGA2/Integrin α2 | HUABIO | Cat# ET1611-57 | 1:1,000 (WB) |
| ITGA3/Integrin α3 | HUABIO | Cat# HA500111; RRID: AB_11156484 | 1:1,000 (WB) |
| ITGAV/Integrin αV | HUABIO | Cat# ET1610-15 | 1:1,000 (WB) 1:300 (ICC) |
| Kif3A | HUABIO | Cat# ER1803-40 | 1:1,000 (WB) |
| KLHL22 | Proteintech | Cat# 16214-1-AP; RRID: AB_2131201 | 1:1,000 (WB) |
| Lamin A/C | Cell Signaling | Cat# 2032; RRID: AB_2136278 | 1:1,000 (WB) |
| LAMP1 | Cell Signaling | Cat# 9091T; RRID: AB_2687579 | 1:1,000 (WB) |
| mCherry | HUABIO | Cat# HA500049 | 1:1,000 (WB) |
| NPC1/Niemann Pick C1 | HUABIO | Cat# ET7107-57 | 1:1,000 (WB) |
| Nrf1/NFE2L1 | Proteintech | Cat# 12936-1-AP; RRID: AB_2267298 | 1:1,000 (WB) |
| PAX6 | Medical & Biological laboratories | Cat# PD022; RRID: AB_ 1520876 | 1:1,000 (IHC) |
| PCNA | Santa Cruz Biotechnology | Cat# SC-56; RRID: AB_628110 | 1:1,000 (WB) |
| PDGFRα | Cell Signaling | Cat# 3174S; RRID: AB_2162345 | 1:1,000 (WB) |
| PD-L1 | Abcam | Cat# Ab282458 | 1:1,000 (WB) |
| Piezo1 | Proteintech | Cat# 15939-1-AP; RRID: AB_2231460 | 1:500 (WB) |
| Profilin-2 | Santa Cruz Biotechnology | Cat# 100955; RRID: AB_2163221 | 1:1,000 (WB) |
| RKIP | Cell Signaling | Cat# 13006S; RRID: AB_2798085 | 1:1,000 (WB) |
| Rpn1/PSMD2 | Bethyl Laboratories | Cat# A303-854A; RRID: AB_2620205 | 1:1,000 (WB) |
| Rpn10/S5a/PSMD4 | Cell Signaling | Cat# 12441S; RRID: AB_2797916 | 1:1,000 (WB) |
| Rpn11/PSMD14 | Cell Signaling | Cat# 4197; RRID: | 1:1,000 (WB) |

| | | | |
|---|---|---|---|
| | | AB_11178935 | |
| Rpn2/PSMD1 | Santa Cruz Biotechnology | Cat# sc-166038 | 1:1,000 (WB) |
| Rpt2/PSMC1 | Proteintech | Cat# 11196-1-AP; RRID: AB_2284521 | 1:1,000 (WB) |
| Rpt3/PSMC4 | Bethyl Laboratories | Cat# A303-849A; RRID: AB_2620200 | 1:1,000 (WB) |
| Rpt6/PSMC5 | Enzo Life Sciences | Cat# BML-PW-9265-0100; RRID: AB_10541436 | 1:1,000 (WB) |
| RRas | HUABIO | Cat# ER60170 | 1:1,000 (WB) |
| SOX2 | Abcam | Cat# Ab79351; RRID: AB_10710406 | 1:200 (IHC) |
| Src | Cell Signaling | Cat# 2109S; RRID: AB_2106059 | 1:1,000 (WB) |
| STIM1 | Abcam | Cat# Ab108994; RRID: AB_10859115 | 1:1,000 (WB) |
| TC-PTP/PTPN2 | R&D Systems | Cat# MAB1930; RRID: AB_2173232 | 1:1,000 (WB) |
| TGF-β Receptor III | Cell Signaling | Cat# 5544S; RRID: AB_10698740 | 1:1,000 (WB) |
| VPS25 | Proteintech | Cat# 15669-1-AP; RRID: AB_2215019 | 1:1,000 (WB) |
| ZO2 | HUABIO | Cat# R1402-2 | 1:1,000 (WB) |
| α5/PSMA5 | Cell Signaling | Cat# 2457S; RRID: AB_823611 | 1:1,000 (WB) |
| α7/PSMA3 | Cell Signaling | Cat# 12446S; RRID: AB_2797918 | 1:1,000 (WB) |
| β5/PSMB5 | Cell Signaling | Cat# 12919S; RRID: AB_ 2798061 | 1:1,000 (WB) |
| β-tubulin | Cell Signaling | Cat# 15115; RRID: AB_2798712 | 1:1,000 (WB) |
| γ-tubulin | HUABIO | Cat# M1701-13 | 1:1,000 (WB) |
| Peroxidase Streptavidin | Jackson Immuno Research | Cat#016-030-084; RRID: AB_2337238 | 1:5,000 (WB) |
| Peroxidase AffiniPure Goat Anti-Rabbit IgG(H+L) | Jackson Immuno Research | Cat# 111-035-003; RRID: AB_2313567 | 1:10,000 (WB) |
| Peroxidase AffiniPure Goat Anti-Mouse IgG(H+L) | Jackson Immuno Research | Cat#115-035-003; RRID: AB_10015289 | 1:10,000 (WB) |
| Peroxidase AffiniPure Goat Anti-Mouse IgG, Light Chain Specific | Jackson Immuno Research | Cat# 115-035-174; RRID: AB_2338512 | 1:10,000 (WB) |

| | | | |
|---|---|---|---|
| Peroxidase Monoclonal Mouse Anti-Rabbit IgG, Light Chain Specific | Jackson Immuno Research | Cat# 211-032-171; RRID: AB_2339149 | 1:10,000 (WB) |
| Alexa 568-goat anti-mouse secondary antibody | Thermo Fisher | Cat# 1841757; RRID: AB_144696 | 1:5,000 (ICC) |
| Alexa 488-goat anti-rabbit secondary antibody | Thermo Fisher | Cat# 1851447; RRID: AB_2576217 | 1:5,000 (ICC) |
| Alexa Fluor™ 568 goat anti-rabbit IgG (H+L) | Thermo Fisher | Cat# 1832035; RRID: AB_10563566 | 1:5,000 (ICC) |
| Alexa Fluor™ 488 goat anti-mouse IgG (H+L) | Thermo Fisher | Cat# 1874804; RRID: AB_2534088 | 1:5,000 (ICC) |

| **Chemicals and Reagents** | **Source** | **Catalog No.** |
|---|---|---|
| Alkynyl Myristic Acid | Click Chemistry Tools | Cat#1164-5 |
| Biotin-LC-Sulfo-NHS | Confluore | Cat# BBBA-8; CAS# 191671-46-2 |
| Biotin-PEG3-Azide | Click Chemistry Tools | Cat# AZ 104-25 |
| Bortezomib | ApexBio | Cat# A2614 |
| BTTAA | Click Chemistry Tools | Cat# 1236-100 |
| Cell Counting Kit-8 | Beyotime | Cat# C0039 |
| Ciprofloxacin | Sigma | Cat# 17850 |
| Collagen Type IV from human placenta | Sigma-Aldrich | Cat# C5533 |
| Cycloheximide | Sigma | Cat# C7698 |
| Digitonin | Sigma | Cat# D141-100MG |
| Fibronectin from bovine plasma | Sigma-Aldrich | Cat# F4759 |
| HisPurTM Ni-NTA Resin | Thermo Fisher | Cat# 25214 |
| Human EGF | Peprotech | Cat# Q99075 |
| Hygromycin B | Sigma | Cat# V900372 |
| IMP-1088 | Cayman Chemical | Cat# HY-112258 |
| IPTG | Amresco | Cat# C0039 |
| Laminin from human placenta | Sigma-Aldrich | Cat# L6274 |
| L-ascorbic acid | Sigma | Cat# A7506 |
| L-Arginine:HCl (13C6; 15N4) | Cambridge Isotope Laboratories, Inc. | Cat# CNLM-539-H-0.1; CAS# 202468-25-5 |
| Lipofectamine$^{TM}$2000 Transfection Regent | Thermo Fisher | Cat# 11668500 |
| L-Lysine:2HCl (13C6) | Cambridge Isotope Laboratories, Inc. | Cat# CLM-2247-H-0.25; CAS# 201740-81-0 |
| Lyso-Tracker (Red) | Beyotime | Cat# C1046 |

| | | |
|---|---|---|
| Medium for SILAC | Thermo Fisher | Cat# 88368 |
| Monensin | SelleckChem | Cat# s2324 |
| MG-132 | SelleckChem | Cat# S2619; CAS# 1211877-36-9 |
| Pierce$^{TM}$ High Capacity Streptavidin Agarose Resin | Thermo Fisher | Cat# 20357 |
| Pierce$^{TM}$ Protein G Agarose | Thermo Fisher | Cat# 20397 |
| Polybrene | Sigma | Cat# TR-1003 |
| Polyethylenimine（PEI） | Polysciences | Cat# 23966-1 |
| Puromycin | Thermo Fisher | Cat# A111380 |
| SYBR$^{@}$ Premix Ex Taq$^{TM}$ II | Takara | Cat# 639676 |
| Trypsin/EDTA | Yeasen | Cat# 40126ES60 |
| Versene | Thermo Fisher | Cat# 15040066 |
| Vitronectin from human plasma | Sigma-Aldrich | Cat# V8379 |

**Plasmids, RNA interference and gene editing**

All cDNAs of human proteasome subunits were originally provided by Dr. Shigeo Murata (The University of Tokyo). CD3δ-GFP was a gift from Dr. Yanfen Liu (ShanghaiTech University). hTERT and N-Ras$^{G12V}$ expression constructs were provided by Drs. Shengda Lin and Bin Zhao, respectively (Zhejiang University). Rpn11-TBHA, Rpt2-internal Flag (IF), Myr$^{Src}$-Rpt2-G2A and Myr$^{Rpt2}$-GFPodc have been reported (*27, 28*). CP8 and uTEV3 were synthesized based on published sequences (*37, 61*). Other cDNAs were PCR-amplified from the ORF LITE human cDNA library (Thermo) or reverse-transcribed from WT MEFs. Insertion of short amino acid stretches such as Myr$^{Src}$ (aa. 1-14 of human Src) and the TEV recognition motif were achieved by annealing and ligation of primers containing the corresponding coding sequences. Site-directed mutagenesis was done with QuikChange® (Agilent), Gibson Assembly® (New England Biolab) or the related CloneExpress II kit (Vazyme), and all constructs have been fully confirmed by Sanger sequencing.

RNAi knockdown and simultaneous cDNA re-expression was achieved using the pLL3.7 lentiviral vector (from Dr. Tyler Jacks, Massachusetts Institute of Technology) as reported (27). For CRISPR/Cas9-mediated Rpt2-G2A knock-in, cells were co-transfected with 2 gRNA sequences (in the Cas9-expressing PX458 vector, Addgene) targeting intronic sequences flanking exon 2 of the human *PSMC1* gene, a donor plasmid with the G2A mutation and homology arms, and the i53 plasmid for enhancing repair efficiency (62). Transfected cells were enriched by flow cytometry. Single clones were screened and confirmed by genomic DNA sequencing and western blot analysis.

All oligonucleotide sequences are listed below.

**PCR Primers**

| | |
|---|---|
| pQCXIP-Rpt2-WT-F | AGCACCGGTACCATGGGTCAAAGTCAGAGTGGTGG |
| pQCXIP-Rpt2-G2A-F | AGCACCGGTACCATGGCTCAAAGTCAGAGTGGTGG |
| pQCXIP-Rpt2-WT-R | AGCCTCGAGTTAGAGATACAGCCCCTCAGGGGTGCC |
| His-SUMO-hRpt2-F | CACAGAGAACAGATTGGTGGAGGTCAAAGTCAGAGTGGTGGTC |
| His-SUMO-hRpt2-R | GAGTGCGGCCGCAAGCTTGTCGACTTAGAGATACAGCCCCTC |
| mLYOX-F | CGGCCGCACCGGTCCTCGAGACCATGCGTTTCGCCTGGGCTG |
| mLYOX-R | ATCCGTTAATTAAGCAATTGGCATACGGTGAAATTGTGCAGCCTG |
| mILEUA-F | GCACCGGTCCTCGAGACCATGGAGCAGCTGAGTTCAGCC |
| mILEUA-R | GCAATTGGCTGGGGAACAAACCCTG |
| mTIMP3-F | CCTCGAGACCATGACTCCCTGGCTTGGGCTT |
| mTIMP3-R | GCAATTGGCGGGGTCTGTGGCGTT |
| hTERT-qPCR-F | GAGCTGCTCAGGTCTTTCTTT |
| hTERT-qPCR-R | CCTCTTCAAGTGCTGTCTGATT |

**Other oligonucleotide sequences**

| | |
|---|---|
| hRpt2-gRNA1-F | CACCGTAAGATCACCCCCCTCTTGA |
| hRpt2-gRNA1-R | AAACTCAAGAGGGGGGTGATCTTAC |
| hRpt2-gRNA2-F | CACCGTCTCGGCTGAGAAGTTCTG |
| hRpt2-gRNA2-R | AAACCAGAACTTCTCAGCCGAGAC |
| shRNA-hRpt2 | GGGGAGTTGCCCAGAGGAA |
| shRNA-mRpt2 | GGAAAATGGTTGGGAGATT |
| TEV-AgeI-F | CCGGCGAGAATCTGTACTTTCAAGGAC |
| TEV-AgeI-R | CCGGGTCCTTGAAAGTACAGATTCTCG |

| | |
|---|---|
| $Myr^{Src}$-NheI-F | CTAGCACCATGGGGAGCAGCAAGAGCAAGCCCAAGGATCCCAGCCAGCGCA |
| $Myr^{Src}$-AgeI-R | CCGGTGCGCTGGCTGGGATCCTTGGGCTTGCTCTTGCTGCTCCCCATGGTG |

**Cell culture, transfection and infection**

WT and G2A MEFs were isolated from E12.5 embryos of the same litter and genotyped. Primary cells were immortalized by infecting with retroviruses expressing SV40 large T/small T antigens (pBabe-T/t) followed by puromycin selection. All other cells were lab stocks originally obtained from American Type Culture Collection (ATCC), with the exception of SH-SY5Y cells which were purchased from National Collection of Authenticated Cell Cultures of China. Cells were maintained in DMEM or RPMI medium supplemented with 10% fetal bovine serum (FBS) and penicillin/streptomycin (all from Thermo). Ciprofloxacin was added periodically to prevent mycoplasma growth. Transfection was done with polyethylenimine (PEI) or Lipofectamine™ 2000 according to standard protocols. Retroviral and lentiviral packaging and infection were performed as previously described (*27*).

**Immunoblotting and immunoprecipitation**

Samples for routine immunoblotting and immunoprecipitation were prepared in the buffer TBSN (50 mM Tris, pH 7.5, 125 mM NaCl, 0.5-1.0% NP-40) supplemented with protease inhibitors (1 mM Pefabloc, 1 mM benzamidine hydrochloride, 1 μM Leupetin, 1 μM E-64 and 1 mM phenylmethanesulfonyl fluoride). When necessary, phosphatase inhibitors were included (10 mM NaF, 20 mM β-glycerolphosphate, 50 nM Okadaic acid and 1-10 mM activated orthovanadate). All procedures were performed on ice or at 4℃. For membrane protein extraction, RIPA buffer (20 mM Tris-HCl, pH 7.4, 150 mM NaCl, 1% Triton X-100, 0.1% SDS, 0.5% sodium deoxycholate) or 1% SDS lysis buffer (in 50 mM Tris, pH 7.5) was used. Protein concentration was determined by Bradford protein assay (Bio-Rad) or BCA protein assay (Thermo). For immunoprecipitation, 0.5-1.0 mg of cell lysate was either incubated with 2-4 μg of antibody for 1 h then with 8-10 μl of Protein G agarose (Thermo) for another 30-45 min, or with 8-10 μl anti-Flag resin for 1 h. Samples were mixed with Laemmli sample buffer, boiled at 95℃ for 5-10 min or incubated at room temperature for 30-60 min (for certain membrane proteins), resolved by SDS-PAGE and transferred to nitrocellulose membranes for immunoblotting.

To assess sterol-induced degradation of HMGCR (*63*), 8.0 x $10^5$ WT or G2A MEFs were seeded in 6 cm plates. On the third day after seeding, cells were sterol-starved for 16 hours by switching into cholesterol-depletion medium (DMEM containing 5% lipoprotein-deficient serum, 1 μM lovastatin and 10 μM mevalonate). Various concentrations of 25-hydroxycholesterol (0/0.1/0.3/1.0 μg/mL) and mevalonate (0/1/3/10 mM) were then added to cells in cholesterol-depletion medium. After 4 hours of treatment, cells were harvested in RIPA buffer containing protease inhibitors (1 mM PMSF, 10 μg/mL leupeptin, 5 μg/mL pepstatin A, 25 μg/mL ALLN and 5 μM MG-132). Cell lysates were centrifuged at 13,000 × g for 10 min and mixed with an equal volume of solubilization buffer (62.5 mM Tris-HCl pH 6.8, 15% SDS, 8 M urea, 10 % glycerol and 100 mM dithiothreitol) and incubated at 37℃ for 30 min. Samples were then mixed with Laemmli sample buffer, resolved by SDS-PAGE and transferred onto polyvinylidene fluoride (PVDF) membrane for immunoblotting.

### Click chemistry and surface biotinylation

Cells were treated with Alk-Myr (50 μM) for 12-24 h and lysed with TBSN buffer as described above. After streptavidin pulldown or anti-Flag immunoprecipitation, the beads were washed and incubated in the click reaction solution: 1 mM biotin-PEG3-azide, 150 μM $CuSO_4$, 300 μM BTTAA and 1 mM TCEP (or 5 mM ascorbic acid) at 30℃ for 1 h. The reaction was stopped by adding 1 mM EDTA and boiled for Streptavidin-HRP blotting.

For surface biotinylation, cells were washed three times with ice-cold PBS then labeled with Biotin-LC-Sulfo-NHS (0.5 mg/mL in PBS, Confluore) at 4℃ for 1 hour with gentle agitation. The reaction was quenched by three 5-min washes with 100 mM glycine in PBS. Cells were lysed with RIPA buffer and sonicated. Spin-cleared cell lysates were then incubated with streptavidin beads at 4℃ for 1 hour. Beads were washed with PBS for 3 times and used for subsequent analysis.

### Proteasome isolation and analyses

Affinity purification of 26S proteasomes using the TBHA-streptavidin system and subsequent elution with TEV cleavage for MS analysis were carried out as described (*23*). Measurement of peptidase activity of the proteasome with the fluorogenic substrate Suc-LLVY-AMC, native gel electrophoresis and sucrose gradient ultracentrifugation were performed as previously reported (*31, 64*). His-SUMO-Rpt2 purification was done as reported (*27*).

**Electron microscopy**

A pre-embedding protocol was used for immunoelectron microscopy (IEM) on brain tissue sections. One-month old male mice (Rpn11$^{TBHA/+}$) were anaesthetized and perfused transcardially with ice-cold PBS (pH 7.4) followed by the fixative (0.1% glutaraldehyde, 4% formaldehyde in 0.1 M phosphate buffer, pH 7.4). Mouse brain was quickly dissected and further fixed overnight. Coronal sections (50 µm) were cut using oscillating microtome (Leica VT1000S) in 0.1 M phosphate buffer (PB, pH 7.4) and post-fixed in the same fixative above for 2 hours. After rinse with PBS (pH 7.4), the brain sections were incubated with 50 mM glycine in 0.1 M PB for 30 min then permeabilized with 0.01% Triton X-100 in 0.1 M PB for 15 min. After blocking with 0.1% BSA-c (Aurion) in 0.1 M PB, the sections were incubated with a rabbit monoclonal anti-HA antibody (Cell Signaling, #3724, 1:100) at 4°C overnight. Following extensive washes, the secondary antibody (Nanogold® Goat anti-rabbit IgG, Nanoprobes, Inc., 1:100) was added. After overnight incubation at 4°C and washing in 0.1 M PB (pH 7.4), the brain sections were again post-fixed with 2.5% glutaraldehyde in 0.1 M PB and treated with the silver enhancement kit (Nanoprobes, Inc.) according to the manufacturer's procedure. Samples were post-stained in 1% $OsO_4$ and 2% uranyl acetate, dehydrated, embedded in epon and ultrathin sections were cut for imaging.

For IEM on MEFs, a post-embedding method was used. Cells grown on sapphire discs were cryo-immobilized by high-pressure freezing (Wohlwend HPF COMPACT 01) and freeze-substituted in 0.1% uranyl acetate and 5% $H_2O$ in acetone, then infiltrated and embedded in LRW resin or HM20 resin according to the instructions. Ultrathin sections (90 nm thick) were mounted on nickel grids coated with formvar film for immunogold staining. Anti-HA (BioLegend, HA.11) or anti-PSMA2 (Cell Signaling, #2455) antibodies were used at 1:10 or 1:20 dilutions, and goat anti-mouse (G7652) or goat anti-rabbit (G7402) secondary antibodies from Sigma were diluted 1:50. Grids with sections were incubated in solutions by floating the section side down on a drop of solution and transferred sequentially from drop to drop to complete the immunostaining. To cover a grid, 5–50 µL drops of solution were placed on a piece of Parafilm in a Petri dish kept in a moist chamber to avoid drying of the solution. A gold enhancemet kit, GoldEnhance-EM plus 2114 (Nanoprobes, Inc.), was used to enlarge the labeled colloidal gold for 3-10 minutes. After rinsing in distilled water, the section was post-stained with 3% uranyl acetate and Sato's lead.

For morphology analysis, cells grown on sapphire discs were cryo-immobilized as above with 0.25 mm-deep HPF carriers as caps and 1-hexadecene as a cryoprotectant. For optimal ultrastructural preservation, samples were transferred to frozen freeze-substitution medium (acetone containing 1% $OsO_4$, 0.1% uranyl acetate and 5% $H_2O$) under liquid nitrogen and placed in an automatic freeze substitutuion system (Leica AFS2, Leica Microsystems) precooled to -90℃. Freeze-substitution was carried out at -90℃ for 8 h. The samples were subsequently warmed to -60℃ over 3 h, kept at -60℃ for 3 h, and warmed to -30℃ over 3 h, kept at -30℃ for 3 h. Then the samples were warmed to 4℃ over 3 h, and kept at 4℃ for 15 min before washing with anhydrous acetone. Samples were then infiltrated and embedded in SPI-Pon 812 resin. 90 nm-thick ultrathin sections were cut with an ultramicrotome (Leica EM UC7, Leica Microsystems), and stained with 3% uranyl acetate in 70% methanol/$H_2O$ for 7 min, followed by Sato's lead for 2 min.

All EM images were acquired on a TECNAI G2 Spirit transmission electron microscope (FEI; Eindhoven, Netherlands) operated at 120 kV. Images with clear membrane/organelle structures were chosen for further analyses.

**Fluorescence microscopy and immunohistochemistry**

Immunofluorescence staining of cells was performed as described in (*65*). Images were taken with a spinning disk confocal microscope (Andor) or laser-scanning microscope (ZEISS LSM 880 with AiryScan) under a 60X or 100X oil lens. LysoTracker labeling was done as instructed by the manufacturer, and live cells were imaged with a DV ELITE microscope (Applied Precision Instruments) under a 60X oil lens. Bright-field images were captured using a Nikon Eclipse TS100 inverted microscope with an Oplenic LCC60-HD camera.

For total internal reflection fluorescence imaging (TIRF), U2OS (parental and G2A knock-in) cells were transfected with mG (membrane-targeted GFP) or Rpn10-GFP. The mG construct has the MGCCFSKT sequence fused to the N-terminus of EGFP (*66*) and marks the plasma membrane. Fixed cells were imaged with an Olympus IX83 microscope using a 100X/1.5 NA oil objective under the TIRF-SIM mode. Penetration depth was set at 200 nm. Structured illumination microscopy (SIM) images were acquired and reconstructed using the Wiener deconvolution algorithm as previously described (*67, 68*). Regular wide-field images were also taken to demonstrate pancellular distribution of the bulk of Rpn10-GFP.

Immunohistochemistry staining of frozen sections of embryonic brain was performed as previously described (*69*). Anti-CD31 and hematoxylin/eosin (H&E) staining of paraffin-embedded embryonic tissues was done according to standard protocols.

**Cell and tissue fractionation**

Two 10 cm plates of cells grown at near confluency were scraped off, pelleted and washed with PBS. Each cell pellet was resuspended in a hypotonic lysis buffer (5 mM HEPES, pH 7.9, 5 mM $MgCl_2$, supplemented with 1 mM ATP and protease inhibitors). Digitonin was also included at a final concentration of 0.005% (w/v) to facilitate cell lysis and centrifugal removal of nuclei. After being swollen on ice for 5 min, cells were broken by passing through a 23G syringe needle for 20 times. A small aliquote of the sample was boiled in SDS loading buffer as the whole cell lysate, while the remainder was centrifuged at 1,400 x g at 4°C for 20 min to remove unbroken cells, nuclei and large cytoskeleton complexes (which all contain proteasomes). The resulting supernatant was further centrifuged at 10,000 x g at 4°C for 10 min. The soluble (cytosolic) fraction was transferred to a new tube, and the membrane pellet was washed twice with the same hypotonic buffer as above. RIPA buffer was added to dissolve membrane proteins from the pellet. After centrifugation at 21,130 x g at 4°C for 10 min, the supernatant was boiled as the membrane fraction.

For isolation of membranes from mouse tissues, different organs were disrupted with a glass-Teflon homogenizer in HEPES buffer (20 mM HEPES, pH 7.4, 0.25 M sucrose, 1 mM EDTA, 1 mM EGTA). Tissue homogenates were cleared twice by centrifugation at 1,000 x g. The supernants were then centrifuged at 15,000 x g, and the resulting supernatants were further spun at 198,000 x g. All centrifugation steps were performed at 4℃ for 10 min. The final membrane pellets containing mostly plasma membrane, microsomal membrane, Golgi and endosomes were dissolved in RIPA buffer or 1X Laemmli sample buffer for western blot analysis.

**Mice**

Rpn11-TBHA knock-in mice (*Psmd14*$^{TBHA}$) were generated at the Transgenic Core of University of California - San Diego and previously reported (*31*). Rpt2-G2A knock-in mice (*Psmc1*$^{G2A}$) were created by CRISPR/Cas9-mediated gene editing at Biocytogen (Beijing, China). All genomic modifications have been confirmed by Southern blot, PCR genotyping and Sanger sequencing. Both strains are on the C57Bl/6 background and have been extensively and periodically back-crossed with WT mice. All animals were housed at the Laboratory Animal Center at Zhejiang University, and all routine husbandry and tumor xenograft studies were in full compliance with policies of the Institutional Animal Core and Use Committee (IACUC) and approved animal protocols. Immunocompromized mice (Nu/Nu) were purchased from Shanghai SLAC Laboratory Animal Co. Ltd (Shanghai, China).

For the tumor xenograft study, 3.0 x 10^6 oncogene-transformed MEF cells were mixed 1:1 (v/v) with Matrigel and injected subcutaneously into the flank of each nude mouse (5-week old females). Tumor volume was monitored regularly with a digital calibar and was calculated by the following formula: $V = 1/2 \times width^2 \times length$.

**Cell adhesion, migration and viability assays**

For cell adhesion assays, purified ECM components were prepared in PBS according to the manufacturer's instruction and used to coat 12-well plates as follows: fibronectin (1 μg/$cm^2$), collagen type IV (0.1 mg/$cm^2$), laminin (1 μg/$cm^2$), and vitronectin (0.1 μg/$cm^2$). Coated plates were air-dried at 37℃ overnight. Cells were dissociated with Versene and resuspended in serum-free medium. After cell counting, 2.0 x 10^5 cells were added to each coated well and allowed to attach at 37℃ for 30 min. Unbound cells were removed by washing twice with PBS, and the remaining cells were fixed with 4% paraformaldehyde and stained with 0.5% crystal violet. After extensive washing with distilled water, air-dried plates were photographed. For each condition, a total of 18 random eye-fields from triplicate wells were imaged, and the numbers of adhered cells were quantified by ImageJ. Alternatively, crystal violet of the stained cells was dissolved with 200 μl ethanol plus 1% (v/v) of concentrated HCl, and Abs590 was measured on a Tecan multiwell plate reader.

For wound-healing (scratch) assays, 8.0 x 10^5 MEFs were seeded in a 12-well plate for 24 hours. A wound through the confluent cell monolayer was made with a P200 micropipette tip. Cells were washed three times with PBS and fresh medium was added. Photos of the wound areas (time "0") were taken. Cells were allowed to migrate for 18 hours, and the same eye-fields were imaged again. Reduction in the wound area was measured by ImageJ and used to reflect cell migration.

For transwell assays, 2.0 x 10^4 MEFs resuspended in serum-free medium were added to the top of a Boyden chamber, while the bottom chamber was filled with DMEM + 10% FBS. Cells were incubated at 37℃ for 24 hours. Migrated cells on the porous membrane were fix-stained with crystal violet as above, imaged and counted.

Cell proliferation and viability were measured by MTS assay or using the CCK-8 kit as previously reported (*31*).

**Quantitative proteomics**

Immortalized MEFs were grown in light or heavy ($^{13}C_6$-Lysine/$^{13}C_6{}^{15}N_4$-Arginine) SILAC medium for more than 7 passages. The cells were washed twice with PBS, harvested in a denaturing lysis buffer (8 M urea, 100 mM Tris, pH 8.5) and sonicated at 4℃ for 10 min. Samples were reduced with 5 mM TCEP (Sigma) at room temperature for 20 min and alkylated with 10 mM iodoacetamide (Sigma) for 15 min in the dark. Urea concentration was diluted to 2 M with 100 mM Tris (pH 8.5) before Trypsin (Promega) was added at a 100:1 protein:enzyme ratio. After digestion at 37℃ for 16 h, the peptides were loaded onto a column which was filled with C18 (3 μm) and SCX (5 μm, Phenomenex) resins and eluted with increasing concentrations of ammonium acetate (25 mM, 75 mM, 100 mM, 150 mM, 200 mM, 275 mM, 375 mM, 500 mM and 1 M). Fractions were vacuum-dried and resuspended in water with 0.1% formic acid. Mass spectrometry experiments were performed on a Q Exactive HF-X instrument (Thermo Fisher Scientific) coupled with an Easy-nLC 1200 liquid chromatography system. Mobile phase A was water, and Mobile phase B was 80% acetonitrile, both containing 0.1% formic acid. Samples were loaded directly onto a C18 reverse phase column (75 μm × 15 cm, 1.9 μm C18). Peptides were separated using a linear gradient of 8–40% B for 50 min, 40–100% B for 5 min, 100% B for 5 min. Data-dependent analysis (DDA) was performed by acquiring a full scan over a m/z range of 350-1500 in the Orbitrap at R = 60,000, NCE = 27, with a normalized AGC target of $3 \times 10^6$. The AGC targets and maximum ion injection time for the MS2 scans were $5 \times 10^4$ and 30 ms, respectively. Precursors of the +1, +8 or above, or unassigned charge states were rejected; exclusion of isotopes was disabled; dynamic exclusion was set to 40 s. Raw data were searched with the MaxQuant software (Version 1.6.10.43) against the mouse database downloaded from UniProt. Peptides were labeled as Arg10 and Lys6; fixed modification was carbamidomethyl; main search peptide tolerance was 4.5 ppm.

**Transcriptomic analysis**

Total RNA was isolated from immortalized MEFs or 3 pairs (WT and G2A littermates) of E12.5 fetal livers using the Trizol method. mRNA-seq was done by BGI (Shenzhen, China). Raw reads were trimmed to 50 bp and mapped to the mouse genome (mm9) using Tophat v2.1.1 with default parameters. Only uniquely mapped reads were subsequently assembled into transcripts guided by the reference annotation (UCSC gene models) using Cufflinks v2.2.1. The RNA abundance of each gene was quantified with FPKM (fragments per kilobase of exon per million mapped fragments). Genes with FPKM < 1 in all samples were excluded, and for the remaining genes, all FPKM values smaller than 1 were set to 1 in subsequent analyses. Differential expressed genes were identified using the criteria |FC| > 2. GO enrichment was performed using "metascape" web server.

#### Data analyses

Western blots were quantified by ImageJ (https://imagej.nih.gov/ij/). Fluorescence microscopy images were processed using the Metamorph® software package (Molecular Devices) or ImageJ. Sequence alignment was done by ClustalW (https://www.genome.jp/tools-bin/clustalw) and ESPript (http://espript.ibcp.fr/ESPript/cgi-bin/ESPript.cgi). Unless otherwise noted, at least three independent experiments were performed to obtain results for statistical analyses (with GraphPad Prism), and quantitative results are shown as mean ± S.D.

Fig. S1

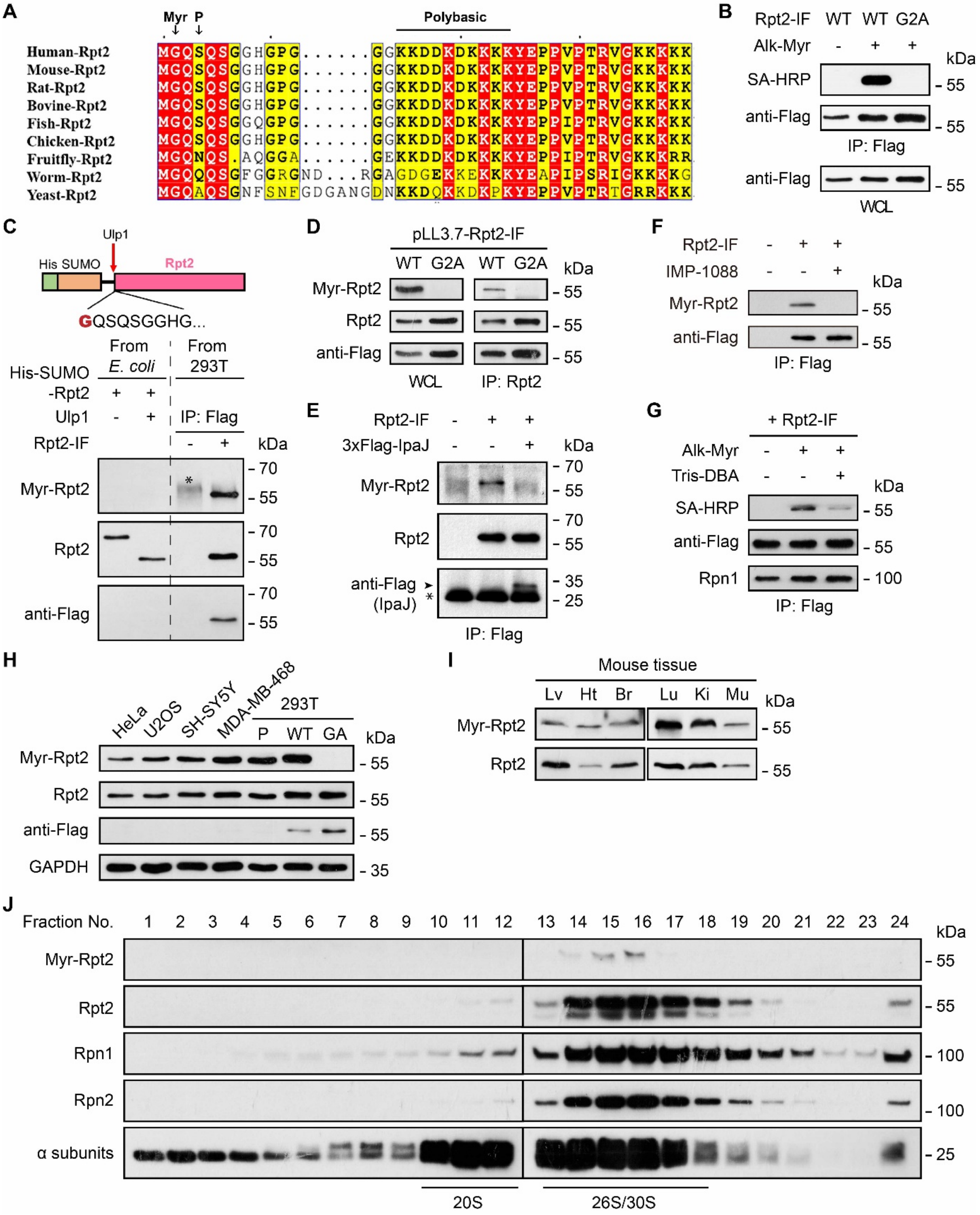


**Fig. S1. Detection and characterization of Rpt2 N-myristoylation.**

(**A**) Alignment of Rpt2 N-terminal sequences from different species. The N-myristoylation site (Gly2), phosphosite Ser4 and the nearby polybasic stretch are indicated.

(**B**) 293T cells were transfected with the indicated Rpt2-IF constructs (“internal Flag”) (*2*) and treated with or without Alk-Myr. After anti-Flag immunopreciptation, Rpt2 myristoylation was detected by click chemistry.
(**C**) Validation of the Rpt2 myristoylation-specific antibody. (Top) A schematic of recombinant His-SUMO-Rpt2 purified from *E. coli*. After Ulp1 cleavage, the His-SUMO tag was removed, leaving a Rpt2 protein with an unmodified N-termimus. (Bottom) Purified His-SUMO-Rpt2 proteins before and after Ulp1 cleavage were probed with the indicated antibodies. WT Rpt2-IF expressed in 293T cells was used as a positive control for Rpt2 myristoylation. Asterisk, a non-specific band.
(**D**) 293T cells were stably transduced with pLL3.7-Rpt2-IF (WT or G2A) for knockdown of endogenous Rpt2 and simultaneous expression of the Rpt2-IF variants. Whole cell lysates and anti-Rpt2 immunoprecipitates were probed with the indicated antibodies.
(**E**) 293T cells were transfected with Rpt2-IF and IpaJ-3XFlag as indicated. Cells were lysed 24 h later for anti-Flag immunoprecipitation and immunoblotting. Arrowhead, Flag-tagged IpaJ. Asterisk, the light chain of the anti-Flag antibody.
(**F**) 293T cells were transfected with or without Rpt2-IF (WT). DMSO or IMP-1088 (1 μM) was added upon transfection. Cells were lysed after 12 h and newly synthesized (transfected) Rpt2-IF was immunoprecipitated and probed with the indicated antibodies.
(**G**) 293T cells were transfected with Rpt2-IF (WT) and immediately treated with Alk-Myr in the absence or presence of the NMT1 inhibitor, Tris-DBA (10 μg/ml), as indicated. Rpt2-IF was immunoprecipitated, click-labeled and probed as in (B).
(**H**) Western blot detection of Rpt2 myristoylation from the indicated cell lines. “WT” and “GA” were 293T cells stably transduced with pLL3.7-Rpt2-IF (WT or G2A) as in (D), used as control.
(**I**) Straight western blot detection of Rpt2 myristoylation from the indicated mouse tissues. Lv, liver; Ht, heart; Br, brain; Lu, lung; Ki, kidney; Mu, muscle.
(**J**) Whole cell lysate of 293T cells was fractionated by sucrose gradient ultracentrifugation. Rpt2 myristoylation and proteasome subunits in each fraction were analyzed by western blot.

**Fig. S2**

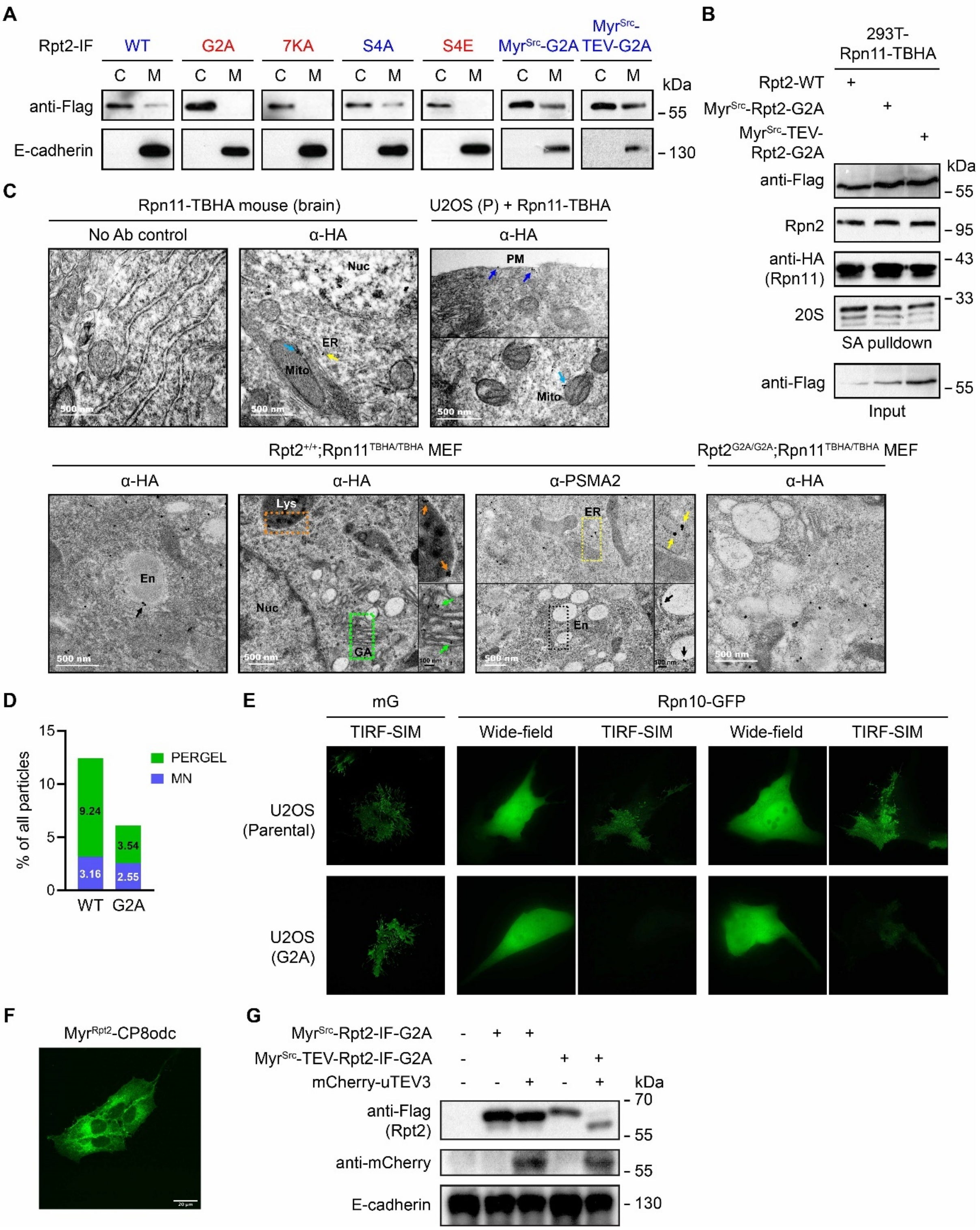


**Fig. S2. The N-terminal sequence of Rpt2 determines proteasome membrane localization.**

(**A**) 293T cells were transfected with the indicated Rpt2-IF variants. After cell fractionation, cytosolic (C) and membrane (M) samples were probed. Rpt2-IF variants capable of membrane-binding were labeled blue, whereas non-membrane-binding mutants were labeled red. 7KA, K15/16/19/21/22/23/24A.

(**B**) 293T-Rpn11-TBHA stable cells were transfected with the indicated Rpt2-IF variants. Their incorporation into the 26S proteasome was confirmed by strepavidin pulldown and anti-Flag immunoblotting.

(**C**) Representative IEM images from the indicated cell/tissue samples stained with antibodies (anti-HA or anti-PSMA2) or without antibodies (as control). Nuc, nucleus; ER, endoplasmic reticulum; Mito, mitochondrion; PM, plasma membrane; En, endosome; Lys, lysosome; GA, Golgi apparatus.

(**D**) Quantification of membrane-associated gold particles (26S proteasomes) in WT and G2A cells. All micrographs from both cell types (MEF and U2OS) with both antibodies (anti-HA and anti-PSMA2) were taken into account. Rpt2-G2A mutation mainly reduced proteasome association with the plasma membrane, ER, Golgi apparatus, endosome/lysosome compartments (“PERGEL”), but less so with regard to proteasomes at the mitochondria or nuclear envelope (“MN”).

(**E**) U2OS (parental and G2A) cells grown on coverslips were transfected with mG or Rpn10-GFP, fixed and imaged with SIM under wide-field or TIRF mode. X = Y = 2,048 pixels, 32.5 nm/pixel.

(**F**) A single clone of U2OS cells stably expressing the $Myr^{Rpt2}$-CP8odc reporter were imaged by a con-focal fluorescence microscope. Scale bar = 20 μm.

(**G**) The indicated $Myr^{Src}$-Rpt2 variants were co-transfected with mCherry-uTEV3 into 293T cells. Cleavage of $Myr^{Src}$-TEV-Rpt2-G2A was confirmed by western blot.

Fig. S3

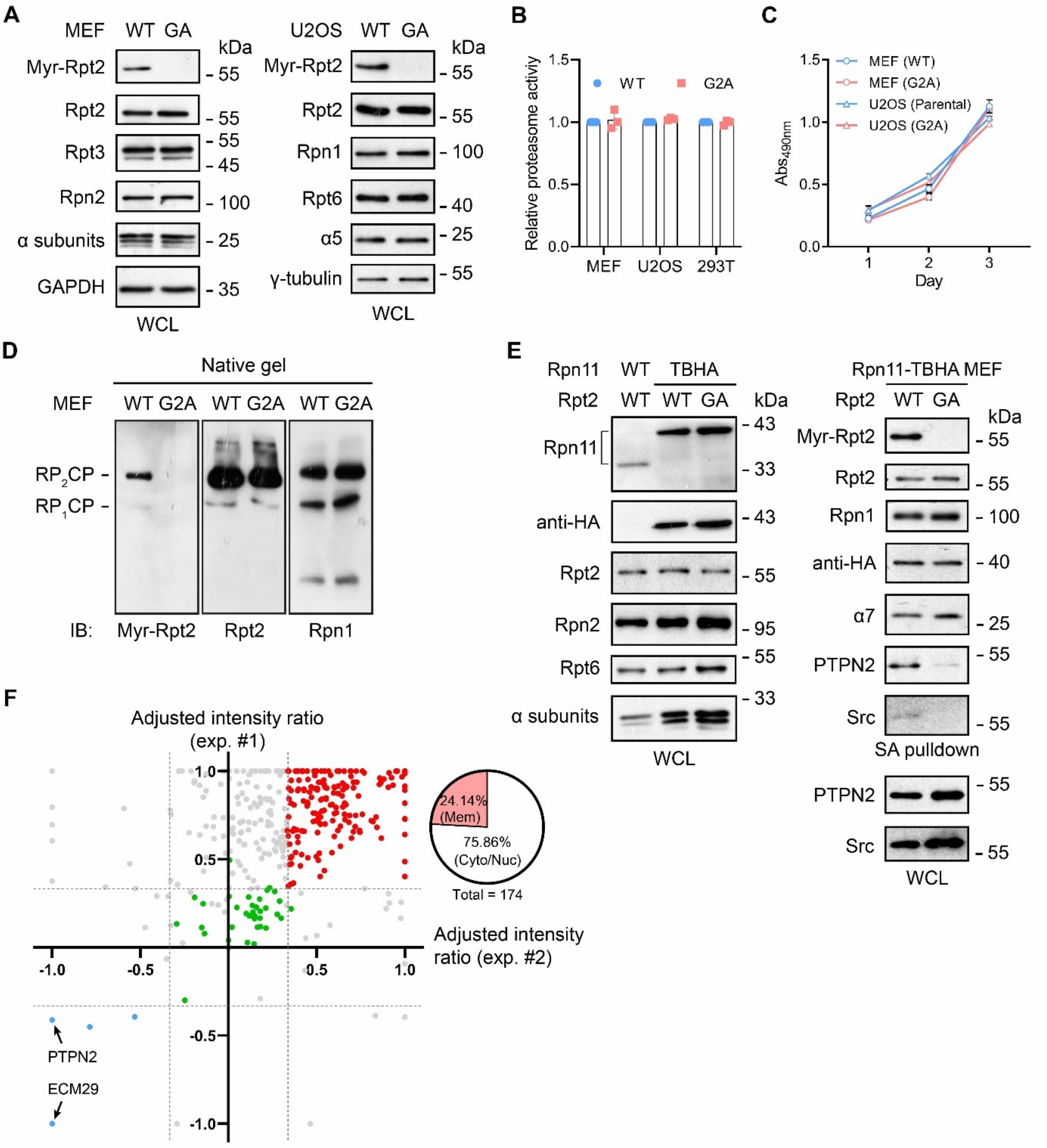


**Fig. S3. Effects of Rpt2-G2A mutation on proteasome assembly, activity, interactome and cell viability.**

(**A**) Western blot analysis of proteasome subunits in WT and G2A cells.

(**B**) Rpt2-G2A knock-in (MEFs, U2OS cells) or replacement (293T pLL3.7 cells) did not affect proteasome peptidase activity against the fluorigenic peptide substrate Suc-LLVY-AMC (n = 3).

(**C**) Viability and proliferation of the indicated cells were measured by the CCK-8 assay (n = 3).

(**D**) Equal amounts of whole cell lysates from WT and G2A/G2A MEFs were resolved by native PAGE and probed with the indicated antibodies.

(**E**) Western blot analysis showing equal abundance (left) and assembly (right) of the proteasome in $Rpt2^{+/+}$;$Rpn11^{TBHA/TBHA}$ and $Rpt2^{G2A/G2A}$;$Rpn11^{TBHA/TBHA}$ MEFs. Proteasome-bound PTPN2 and Src were also probed.

(**F**) Rpt2-G2A mutation alters the proteasome interactome. Endogenous proteasomes were streptavidin-purified from $Rpt2^{+/+}$;$Rpn11^{TBHA/TBHA}$ and $Rpt2^{G2A/G2A}$; $Rpn11^{TBHA/TBHA}$ MEFs and analyzed by label-free mass spectrometry. Proteasome-interacting proteins (PIPs) identified from two independent experiments were plotted by their adjusted intensity ratios between WT and G2A cells. The adjusted intensity ratio was calculated as $(Intensity_{G2A} – Intensity_{WT})/(Intensity_{G2A} + Intensity_{WT})$, such that a two-fold increase or decrease in G2A cells corresponds a ratio of 0.33 or -0.33, respectively (marked by dotted lines). By this standard, a total of 174 PIPs consistently showed increased proteasome binding in G2A cells (red), over 75% of which were cytosolic or nuclear (non-membrane-associated) proteins. Four PIPs showed decreased binding (blue), including PTPN2 and Ecm29 (arrows). Core proteasome subunits (green) remained essentially unchanged. Other PIPs are labeled gray.

**Fig. S4**

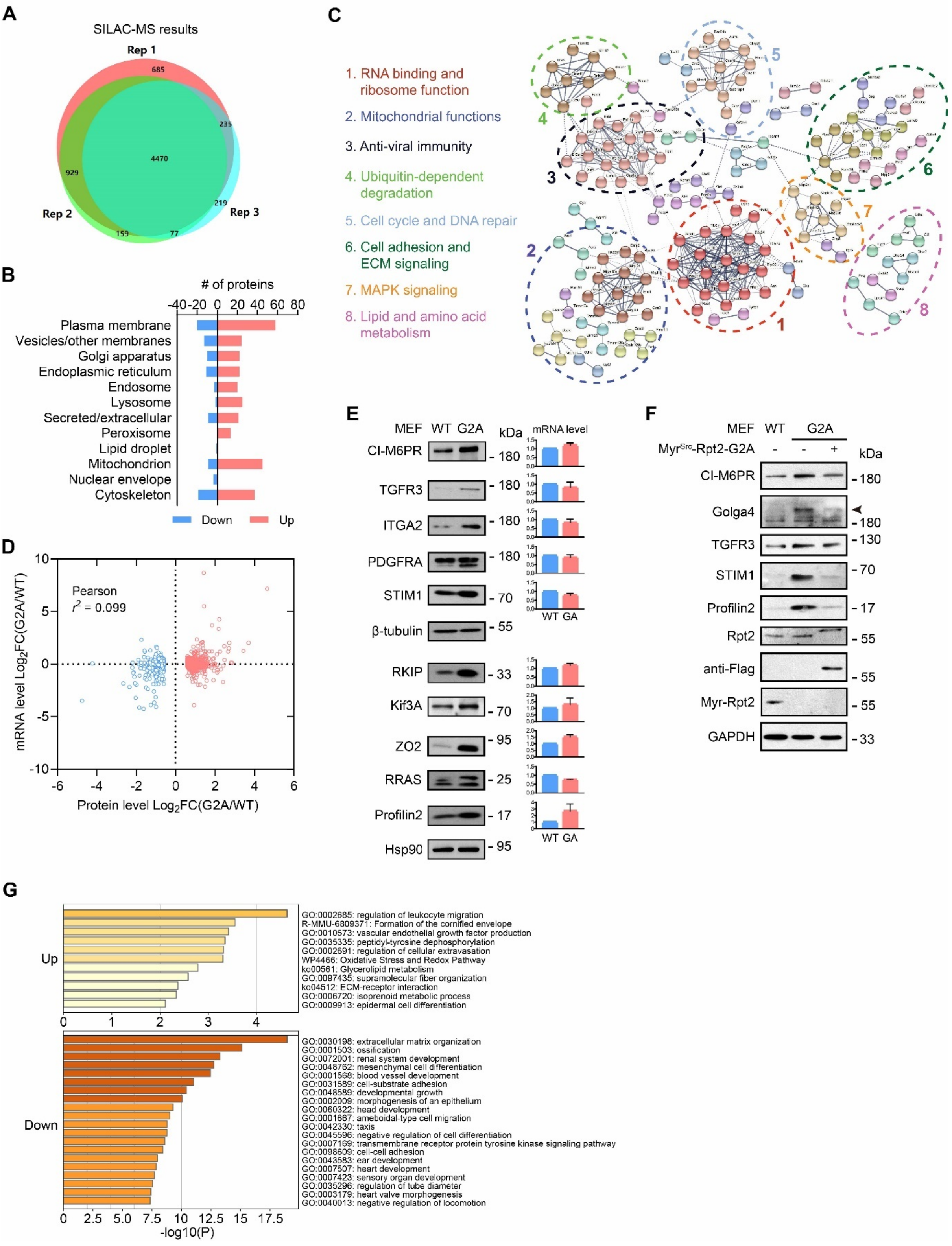


**Fig. S4. Proteomic and transcriptomic changes in Rpt2-G2A cells.**
(**A**) Venn diagram showing the numbers of proteins identified from WT and G2A MEFs in three SILAC-MS experiments.
(**B**) Subcellular distribution of proteins up- and down-regulated in G2A MEFs.

(**C**) Protein-protein interaction network of downregulated proteins in G2A MEFs.
(**D**) Correlation analysis between the protein and mRNA levels of upregulated (red) and downregulated (blue) proteins in G2A cells, based on SILAC-MS and RNA-Seq data.
(**E**) Western blot verification of upregulated transmembrane proteins (left) and peripheral membrane proteins (right) that were identified by SILAC-MS. The corresponding mRNA levels from RNA-Seq analysis are shown on the side.
(**F**) $Myr^{Src}$-Rpt2-G2A expression in G2A MEFs restored the levels of the indicated proteins.
(**G**) GO term analysis of differentially expressed mRNAs identified in G2A MEFs by RNA-Seq.

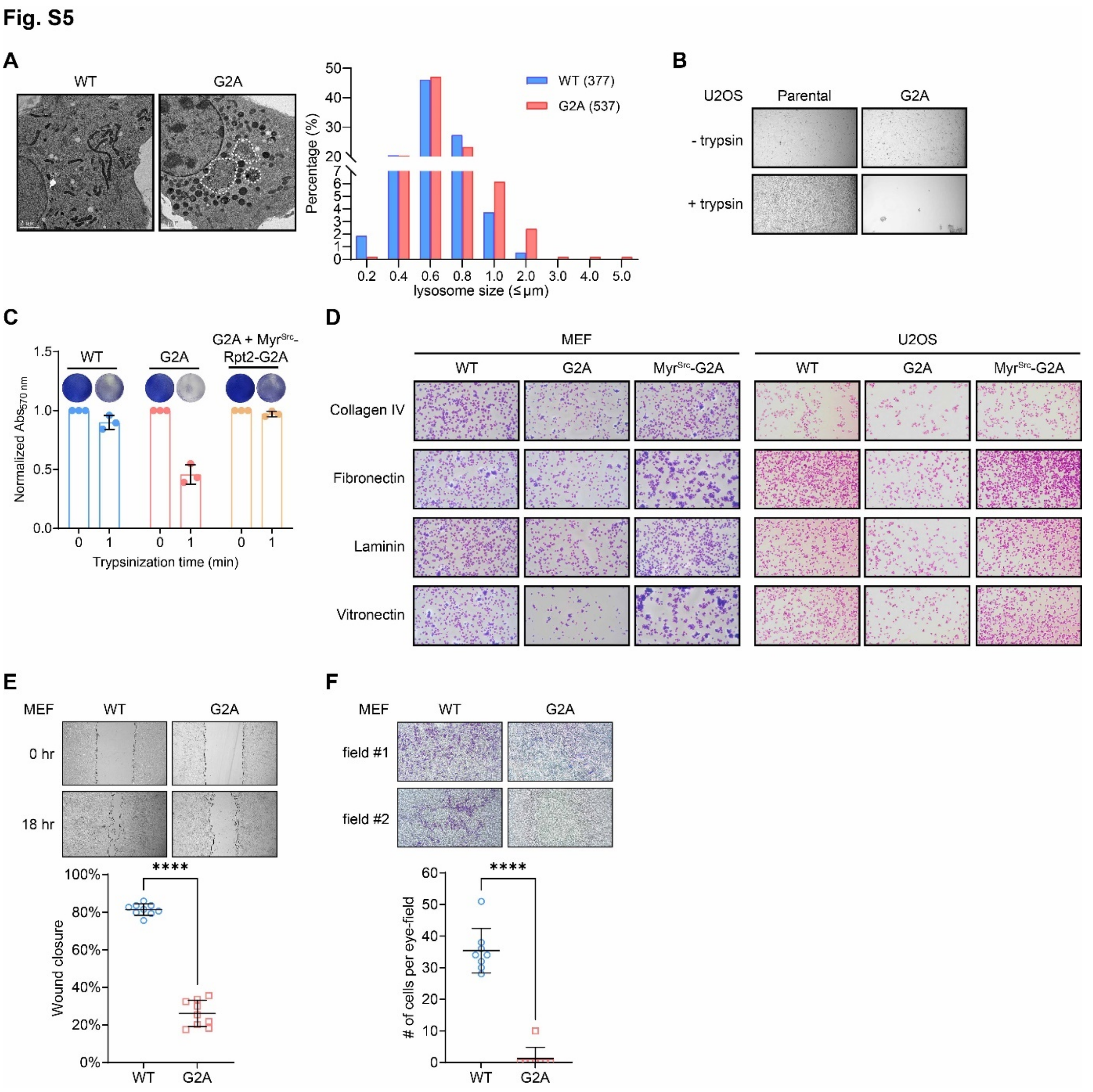


**Fig. S5. Loss of Rpt2 myristoylation undermines cell adhesion, migration and endomembrane homeostasis.**

(**A**) Representative TEM images showing lysosomes/MVBs in WT and G2A MEFs (left). Lysosome sizes were measured and quantified (right). Abnormally large, membrane-bound vacuoles (dotted lines) were occasionally observed in G2A cells but not in WT cells.

(**B**) U2OS cells were seeded in a 12-well plate at 1.0 x $10^5$/well. The next day, cells were washed with PBS and briefly treated with 0.05% trypsin/EDTA at 37°C for 2 min. Photographs were taken before and after trypsinization (objective magnification = 10X).

(**C**) The indicated MEFs were treated without or with trypsin/EDTA (0.0625%, 1 min). Cells were fix-stained with crystal violet immediately after 1-min treatment of trypsin and imaged. After extensive washes, the amounts of crystal violet retained were quantified by measuring absorbance at 570 nm.

(**D**) Representative photographs of crystal violet-stained cells adhered to ECM protein-coated surfaces (objective magnification = 10X).

(**E**) WT and G2A MEFs were allowed to migrate for 18 hours in a scratch/wound-healing assay. Reduction of the wound area was measured and plotted. Objective magnification = 4X. ****, $P < 0.0001$ ($N = 9$ eye-fields from 3 independent experiments, Student's *t*-test).

(**F**) Representative images of porous membranes of the transwell migration assay, with migrated MEFs stained with crytal violet (objective magnification = 10X). The numbers of migrated cells in each eye-field were counted and plotted. ****, $P < 0.0001$ ($N = 8$, Student's *t*-test).

**Fig. S6**

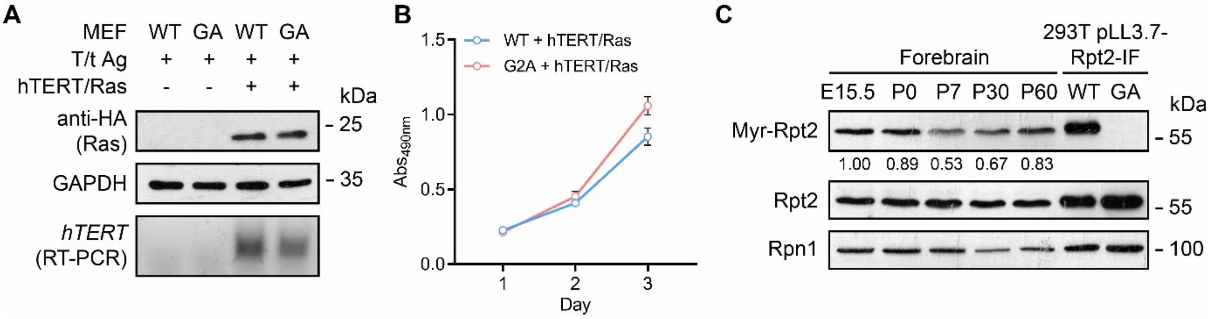


**Fig. S6. Oncogene expression in MEFs and Rpt2 myristoylation during brain development**

(**A**) Stable expression of N-Ras$^{G12V}$ and hTERT in immortalized MEFs was confirmed by western blot (top) and RT-PCR (bottom), respectively.

(**B**) Proliferation of oncogene-transformed MEFs was measured by the CCK-8 assay ($N$ = 3).

(**C**) The dynamics of Rpt2 N-myristoylation level in rodent forebrain from embryonic stage to adulthood was determined by western blot. The ratios between Myr-Rpt2 and total Rpt2 at each developmental stage are shown.
